# Identification of touch neurons underlying dopaminergic pleasurable touch and sexual receptivity

**DOI:** 10.1101/2021.09.22.461355

**Authors:** Leah J. Elias, Melanie Schaffler, Isabella Succi, William Foster, Mark Gradwell, Manon Bohic, Lindsay Ejoh, Victoria Abraira, Ishmail Abdus-Saboor

## Abstract

Pleasurable touch during social behavior is the key to building familial bonds and meaningful connections. One form of social touch occurs during sexual encounters. Although sexual behavior is initiated in part by touch, and touch is ongoing throughout copulation, the identity and role of sensory neurons that transduce sexual touch remain unknown. A population of sensory neurons labeled by the G-protein coupled receptor Mrgprb4 detect stroking touch in mice, however, these neurons have never been implicated in any natural social behaviors. Here, we study the social relevance of Mrgprb4-lineage neurons by genetically engineering mice to allow activation or ablation of this population and reveal that these neurons are required for sexual receptivity and sufficient to induce dopamine release in the brain. Even in social isolation, optogenetic stimulation of Mrgprb4-lineage neurons through the back skin is sufficient to induce a conditioned place preference and a striking dorsiflexion resembling the lordotic copulatory posture in females. In the absence of Mrgprb4-lineage neurons, female mice no longer find male mounts rewarding: sexual receptivity is supplanted by aggression and a coincident decline in dopaminergic release in the mesolimbic reward pathway. In addition to sexual behavior, Mrgprb4-lineage neurons are also required for social postures induced by female-to-female back touch. Together, these findings establish that Mrgprb4-lineage neurons are the first neurons of a skin-to-brain circuit encoding the rewarding quality of social touch.

## Introduction

The pleasure of a partner’s caress or a child’s embrace begins with mechanical signals transduced by neurons in our skin. After detection of touch in the skin by receptor proteins expressed on the surface of dorsal root ganglion (DRG) sensory neurons, electrical signals are transduced to defined neurons in the spinal cord (Jenkins and Lumpkin, 2017). The spinal cord then serves as a local processing hub for somatosensory information before signals reach the brain – the ultimate source for sensory perception (Abraira et al., 2017; Haring et al., 2018). The way our brain interprets instances of social touch is critical for our well-being. Despite the centrality of socially rewarding touch in our daily lives, the neurons in the skin that detect social touch and shape the valence of perception generated in the brain, remain unknown. This gap in knowledge is critical, especially when considering the nature of neurodevelopmental disorders like autism spectrum disorder, where gentle touch and socially rewarding behaviors are aversive (Orefice et al., 2019; Orefice et al., 2016; Peled-Avron and Shamay-Tsoory, 2017).

How might touch generate reward during skin-brain interactions? The mesolimbic pathway in the midbrain consists of ventral tegmental area (VTA) dopaminergic neurons that release dopamine into the nucleus accumbens (NAc) to promote reward-learning and reinforcement, motivation for rewards, and reward-prediction errors that teach animals to alter behavior whenever reward values do not match predictions (Hart et al., 2014; Mohebi et al., 2019; Schultz, 2016; Starkweather et al., 2017; Wise, 2004). Regarding social behaviors in particular, such as exploring a non-familiar conspecific, same-sex social interactions, or play behavior between female rats, VTA dopamine signaling drives these social interactions (Bariselli et al., 2018; Gunaydin et al., 2014; Northcutt and Nguyen, 2014). Although a role for VTA dopamine neurons in promoting social behaviors has been identified, except for a handful of studies (Wang et al., 2021), the neurons and circuits that encode social stimuli and activate VTA dopamine neurons remains obscure. Moreover, considering that dopamine neurons in the VTA are themselves a heterogeneous population (Morales and Margolis, 2017), this raises the possibility that some VTA dopamine neurons might be fine-tuned for the reward associated with touch during behaviors ranging from sex to social bonding.

A class of sensory neurons in humans that are linked to gentle stroking are termed C-tactile afferents (Ackerley et al., 2014; Pawling et al., 2017), and there are putative populations of these neurons in the mouse (Delfini et al., 2013; Handler and Ginty, 2021; Lou et al., 2013). One such class of neurons in mice express the G-protein coupled receptor Mrgprb4 and appear to share anatomical and physiological similarities with human C-tactile afferents (Liu et al., 2007; Vrontou et al., 2013). Prior studies demonstrate that hairy skin-innervating C-fibers labeled in the *Mrgprb4*^*Cre*^ mouse line respond to gentle stroking and produce a conditioned place preference, suggesting that their activation is rewarding (Liu et al., 2007; Vrontou et al., 2013). These findings prompted us to ask whether sensory neurons marked by the *Mrgprb4*^*Cre*^ mouse line might be important for promoting ethologically relevant rewarding touch that engages the mesolimbic reward pathway in the brain. Here we used a combination of mouse genetics, novel behavioral paradigms, and *in vivo* brain imaging to connect the skin and brain by dissecting the role of Mrgprb4-lineage neurons in socially rewarding behaviors.

## Results

### Focalized activation of Mrgprb4-lineage neurons in the skin is rewarding and induces lordosis-like posture in female mice

We reasoned that focal activation of Mrgprb4-lineage neurons in the back skin might mimic touch among mice, so we used the blue light sensitive ion channel channelrhodopsin (ChR2) to focally stimulate Mrgprb4+ neurons *in vivo*. To accomplish this goal, we used an *Mrgprb4*^*Cre*^ driver that functions as a lineage tracer to express a ChR2-eYFP fusion protein in Mrgprb4-lineage neurons (Fig 1a). We confirmed this genetic targeting strategy with RNAscope *in situ* hybridization and immunostaining (Fig 1b,c). Because Mrgprb4 expression begins ∼P4-5 in a population of progenitors broader than the Mrgprb4-expressing population of adult neurons(Liu et al., 2008), we characterized the expression of ChR2-eYFP in Mrgpra3+, Mrgprc11+, and Mrgprd+ populations of dorsal root ganglion neurons, which are known to share lineage with Mrgprb4+ neurons. Using RNAscope *in situ* hybridization to characterize co-expression of ChR2-eYFP with Mrgprb4, Mrgprd, Mrgprc11, and Mrgpra3. (Supplemental Figure 1), we found the expression of ChR2 across these populations consistent with the expected developmental expression of Mrgprb4(Liu et al., 2008) (Fig 1d,e), thus accounting for all ChR2-eYFP+ cells in *Mrgprb4*^*Cre*^*;Rosa*^*ChR2*^ mice. Moreover, there was little to no overlap between ChR2 (Mrgprb4-lineage expression) with TH, which is a marker of other C-LTMRs (Li et al., 2011), confirming that the Mrgprb4 lineage neurons are a distinct population of C-fibers.

**Figure 1:**
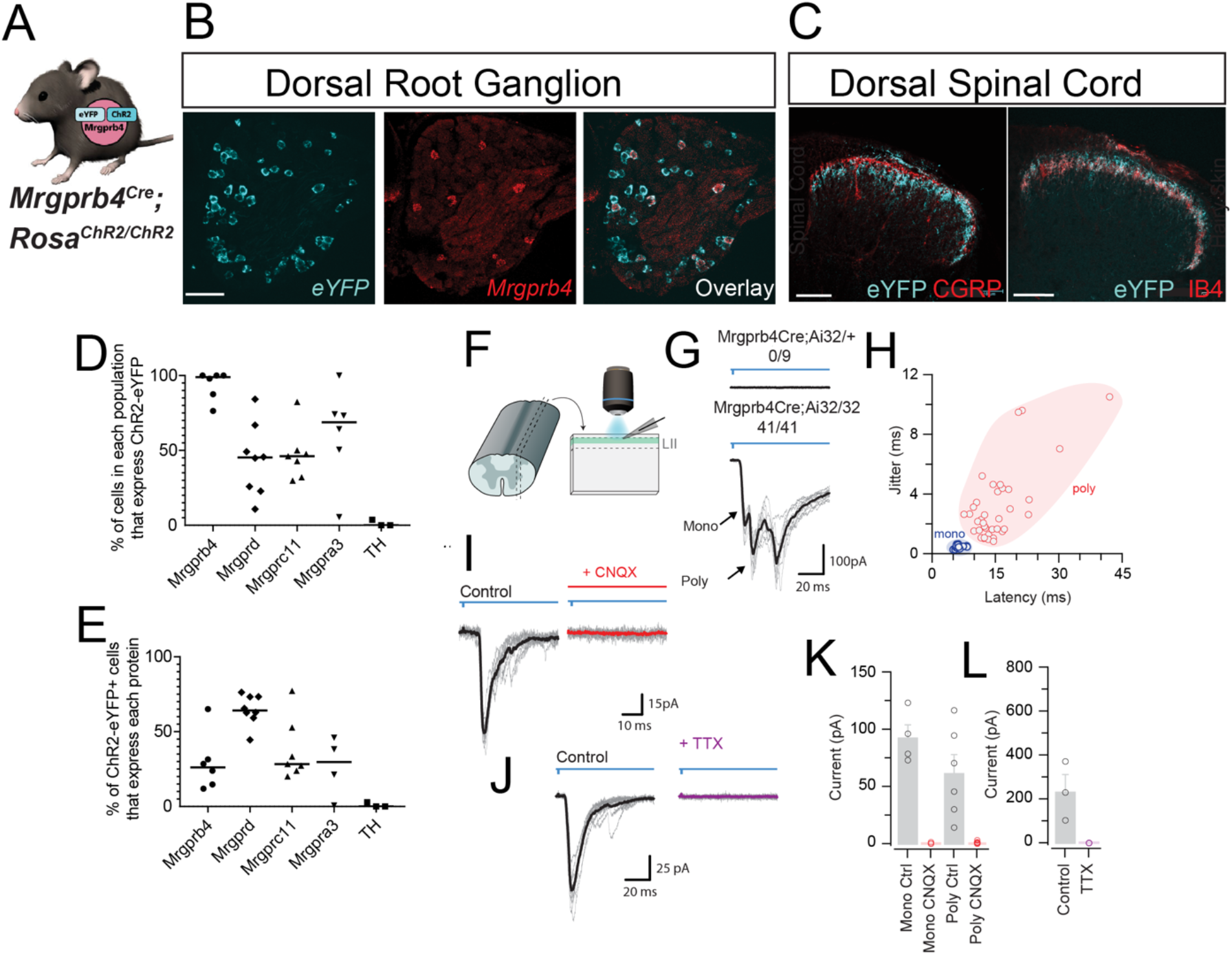
Genetic targeting strategy to optogenetically manipulate Mrgprb4-lineage touch neurons. A) *Mrgprb4*^*Cre*^ mice express ChR2-eYFP in a Cre-dependent manner. B) RNAscope *in situ* hybridization in DRG to quantify the expression of ChR2 in *Mrgprb4+* cells. Scale bars represent 100 µm. C) Immunofluorescence staining in spinal cord dorsal horn. Mrgprb4+ terminals innervate lamina II, inferior to CGRP+ terminals in lamina I and overlapping with IB4+ terminals in lamina II. Scale bars represent 100 µm. D) Expression of *eYFP* in different populations of DRG neurons that share lineage with *Mrgprb4*. From RNAscope *in situ* hybridization in supplemental figure 1. E) Expression of different lineage populations among *ChR2-eYFP*+ cells. See supplemental figure 1 for original RNAscope *in situ* hybridization images. F) Schematic illustrating spinal cord slice electrophysiological recordings from lamina II during optogenetic stimulation of terminals. G) Mono-or poly-synaptic light induced currents only in *Mrgprb4*^*Cre*^; *Rosa*^*ChR2/ChR2*^ mice. H) Quantification of light induced currents from *Mrgprb4*^*Cre*^; *Rosa*^*ChR2/ChR2*^ mice with jitter used to determine mono- or poly-synaptic transmission. I) Light-induced currents in *Mrgprb4*^*Cre*^; *Rosa*^*ChR2/ChR2*^ mice are abolished with CNQX. J) Light-induced currents are abolished in *Mrgprb4*^*Cre*^; *Rosa*^*ChR2/ChR2*^ mice with TTX. K,L) Quantification of data presented in I,J.

With confirmation of our genetic targeting strategy to activate Mrgprb4-lineage neurons with ChR2, we next asked if we could induce light-evoked, synaptically driven currents to second-order neurons downstream in the dorsal horn of the spinal cord by combining optogenetics with whole cell patch clamp physiology (Fig 1f). Interestingly, recording from LII of the spinal cord of a *Mrgprb4*^*Cre*^*;Rosa*^*ChR2/+*^ heterozygous mice yielded no light-induced currents, while all recordings from *Mrgprb4*^*Cre*^*;Rosa*^*ChR2/ChR2*^ homozygotes exhibited mono-and/or poly-synaptic inputs (Fig 1g,h; Supplemental Figure 2). We previously published that two copies of Rosa-ChR2 were needed to evoke behavior in *Mrgprd*^*Cre-ERT2*^*;Rosa*^*ChR2/ChR2*^ mice but not *Trpv1*^*Cre*^*;Rosa*^*ChR2/+*^mice, potentially pointing towards unique and unexplored physiological characteristics of the Mrg lineage of C-fibers^13,14^. 15/39 neurons recorded fired action potentials following photostimulation. Action potentials induced by these inputs were abolished by TTX, and both mono and polysynaptic Mrgprb4 inputs to second-order neurons can be abolished by application of CNQX, demonstrating both sodium channel-dependent firing of Mrgprb4-lineage neurons and glutamatergic synaptic transmission between Mrgprb4-lineage neurons and second-order neurons (Fig 1i-l). Together, the genetic targeting and electrophysiological recordings demonstrate the feasibility of optogenetically activating Mrgprb4-lineage neurons to evoke behavior.

Previous work demonstrated that chemogenetic activation of neurons labeled in the *Mrgprb4*^*Cre*^ mouse line in juvenile males induces a conditioned place preference(Vrontou et al., 2013), suggesting activation of these neurons is inherently pleasant or rewarding. To test whether focal activation of these afferents in the back skin is similarly preferable, we developed a conditioned place preference assay in which each chamber was paired with laser light administered transdermally through the shaved back skin. We used *Mrgprb4*^*Cre*^; *Rosa*^*ChR2/ChR2*^ mice given our result with spinal cord slice physiology regarding the importance of homozygosity of the opsin protein (Fig 1f,g). One chamber had ChR2-activating blue light, and the other had non-stimulating green light as a control (Fig 2a). *Mrgprb4*^*Cre*^; *Rosa*^*ChR2/ChR2*^ females had a significant preference for the chamber associated with blue light stimulation after training compared to Cre-negative littermates, suggesting that focal activation of Mrgprb4-lineage neurons in the back skin is positively reinforcing and inherently rewarding (Fig 2b,c). Because approximately 60% of Mrgprb4-lineage neurons express Mrgprd in adulthood (Fig 1e), we sought to confirm the specificity of this result to Mrgprb4-lineage neurons by repeating these experiments with *Mrgprd*^*Cre-ERT2*^; *Rosa*^*ChR2/ChR2*^ mice. The *Mrgprd*^*Cre-ERT2*^ mouse line has an inducible Cre, allowing us to treat these mice with tamoxifen at weaning, thereby labeling only the adult Mrgprd+ population that does not co-express Mrgprb4. We and others have previously demonstrated that optogenetic activation of Mrgprd+ neurons is neutral at baseline conditions(Abdus-Saboor et al., 2019; Beaudry et al., 2017; Warwick et al., 2021), and we recapitulated those findings here in our newly developed optogenetic placed preference paradigm by showing no preference for blue light when Mrgprd+ neurons are activated through the back skin (Fig 2d,e). Thus, the positive valence is uniquely associated with activation of Mrgprb4-lineage neurons. Focal activation in the back is sufficient to drive this preference, suggesting the Mrgprb4-lineage neurons may detect pleasant tactile contact to the back skin.

**Figure 2:**
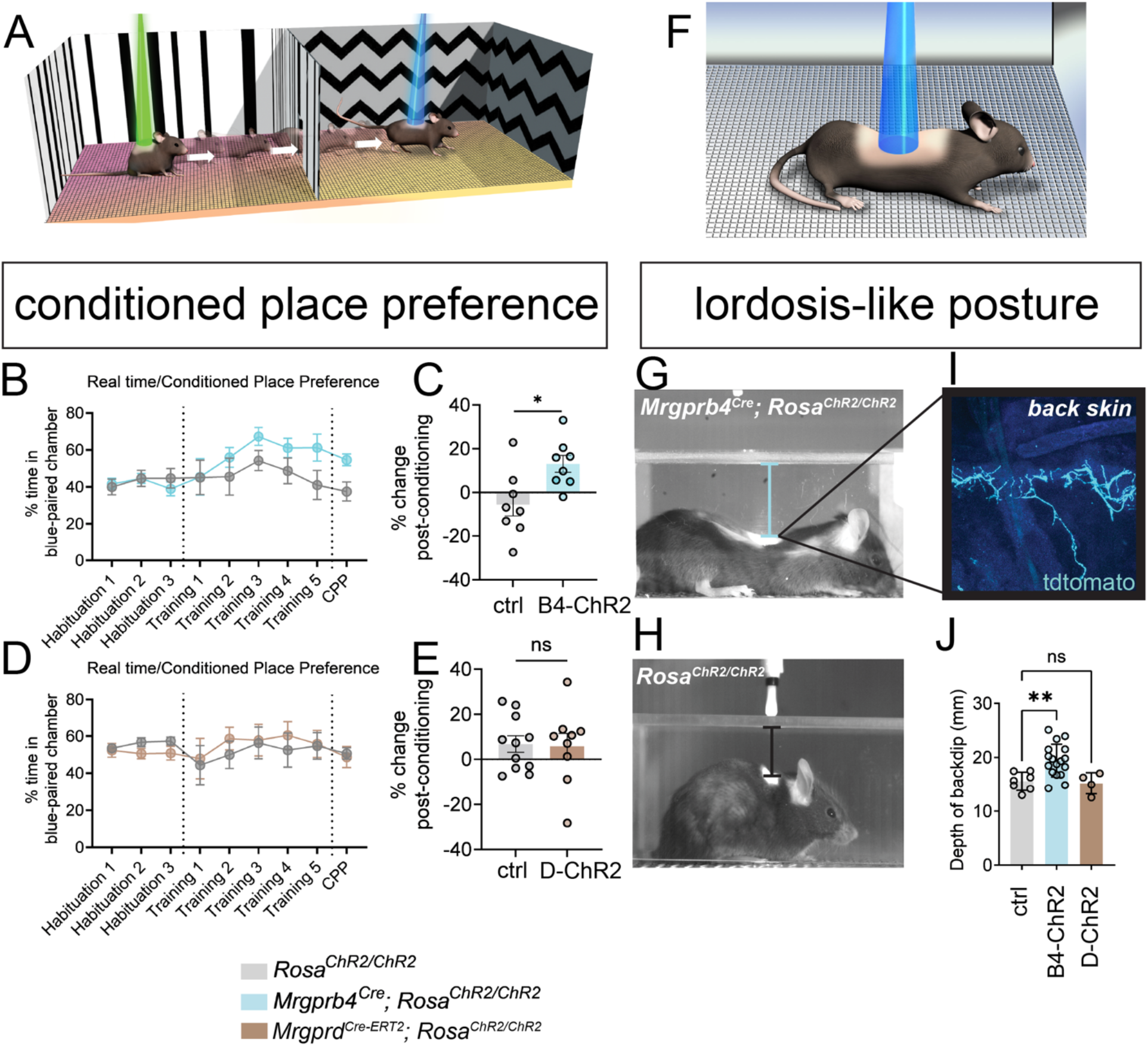
Focalized activation of Mrgprb4-lineage neurons in the back skin is rewarding and induces a lordosis-like posture in female mice. A) Schematic illustrating our real time/conditioned place preference assay. B) *Mrgprb4*^*Cre*^; *Rosa*^*ChR2/ChR2*^ females gradually spend more time in the blue laser chamber compared to green laser chamber during the training days (lasers on) and C) spend significantly more time in the chamber they learned to associate with blue light on the test day (lasers off) compared to habituation days, (\**P<0*.*05*, unpaired t-test.) D,E) *Mrgprd*^*CRE-ERT2*^; *Rosa*^*ChR2/ChR2*^ females do not develop a preference for the blue laser-paired chamber. F,H) Stills from high speed videography to closely examine the behavior during the preferable transdermal optogenetic activation of Mrgprb4-lineage neurons. G) On the right, immunofluorescence staining of Cre-tdtomato+ terminals in the hairy skin. *Mrgprb4*^*Cre*^; *Rosa*^*ChR2/ChR2*^ females (F) exhibit a striking lordosis-like dorsiflexion in response to Mrgprb4-lineage neuron activation, a behavior absent from (H) *Rosa*^*ChR2/ChR2*^ female littermates. I) This posture is quantified as the back’s maximum distance from the chamber ceiling over the course of 20s optogenetic stimulation.

Behaviors that result from optogenetic activation of sensory neurons that sense pain or itch are intuitive to interpret. For example, we and others have shown that selective optogenetic activation of nociceptive neurons in mice evokes pain behaviors, such as paw withdrawal, licking, and shaking, while activation of itch-sensing neurons evokes stereotyped itch behaviors, such as scratching(Abdus-Saboor et al., 2019; Daou et al., 2016; Daou et al., 2013; Iyer et al., 2014; Olson et al., 2017; Sharif et al., 2020; Sun et al., 2017). To uncover a stereotyped behavior evoked by optogenetic activation of Mrgprb4-lineage neurons in freely behaving mice, we combined optogenetic stimulation with high-speed videography at 750 frames per second to capture behavior at high spatial and temporal resolution (Fig 2f). Intriguingly, a lordosis-like posture, involving robust dorsiflexion, appeared as a stereotyped response to selective activation of Mrgprb4-lineage neurons in the back skin (Fig 2g,h). Immunofluorescence staining for tdTomato confirmed that the Mrgprb4-Cre-tdTomato was indeed active in the terminals of the hairy skin of the back, which received the blue light stimulation (Fig 2i). We quantified this posture as the maximum distance the back concaved from the ceiling of the plastic chamber over the course of 20s of optogenetic stimulation (Fig 2j). The posture is not observed upon optogenetic activation in the same experimental chambers of either Mrgprd+ neurons (Fig 2h) or Mrgpra3+ neurons (Supplemental Figure 3), two related populations of neurons that share lineage with Mrgrprb4. Thus, this posture is specific to activation of Mrgprb4-lineage neurons and represents the first stereotyped behavioral response to the selective activation of pleasant touch. Because this response resembles the sexual receptivity lordosis posture and occurred in female mice with optogenetic stimulation directed to the back, we hypothesized that Mrgprb4-lineage neurons might play a role in sexual receptivity and other touch-dependent socially rewarding behaviors.

### Mrgprb4-lineage neurons are required for sexual receptivity and female-female social postures but not social interest

Next, we interrogated the role of Mrgprb4 neurons in two natural behaviors where female rodents display prominent dorsiflexion postures in response to touch: 1) lordosis – the female sexually receptive posture which includes dorsiflexion in response to male tactile input to the back and flanks(Kow et al., 1979; Larsson et al., 1976; Pfaff and Sakuma, 1979a, b; Sodersten and Larsson, 1975) and 2) crawling underneath cage mates which involves a dorsiflexion of the spine in response to the cage mate on top, and has recently been associated with oxytocin neuron activation in rats(Tang et al., 2020). We refer to this behavior as the “conspecific crawl.” We focused on female social behaviors because lordosis is a female behavior and conspecific tactile input may be conflated with aggression or dominance behavior in males.

To determine whether Mrgprb4-lineage neurons are required for either of these touch-dependent social behaviors, we ablated the population using a Cre-dependent diphtheria toxin mediated ablation (Fig 3a; lineage quantification in Fig 1d,e). We used RNAscope *in situ* hybridization to confirm the elimination of dorsal root ganglion (DRG) cell bodies that express *Mrgprb4* (Fig 3b). The same population of Mrgprb4-lineage neurons were ablated upon DTA expression as were labeled with ChR2-eYFP expression (this RNAscope data to be published in a separate study). *Mrgprb4*^*Cre*^; *Rosa*^*DTA*^ mice did not exhibit any motor abnormalities that might conflate interpretations of our results (Fig 3c,d). To determine if the Mrgprb4-lineage neurons are required in females to detect male mounts and facilitate lordosis, we conducted a lordosis quotient assay(McCarthy et al., 2017; Pfaff and Sakuma, 1979b). To control for natural fluctuations in the estrous cycle, we ovariectomized females (OVX) and subsequently treated them with both estradiol and progesterone to mimic a state of behavioral estrus at the time of the assay, as previously described(Frye et al., 2014) (Fig 3e). *Mrgprb4*^*Cre*^; *Rosa*^*DTA*^ females and cage mate controls were paired with stud males that had recently fathered multiple litters, confirming their sexual competency. We measured the fraction of receptive responses to total male mounts (lordosis quotient) as well as the average duration the female maintained a receptive posture (sexual receptivity). A female’s response was considered receptive if she had all four limbs on the floor of the cage and displayed no attempts to escape. *Mrgprb4*^*Cre*^; *Rosa*^*DTA*^ females had similar lordosis quotient and sexual receptivity to controls on the first week of testing, but, strikingly, their receptivity plummeted on the second pairing, and remained minimal for the third pairing (Fig 3g,h,k,l). On the other hand, control females increased in receptivity with each trial, suggesting that intact female mice might be learning to engage in this rewarding behavior (Fig 3g,h).

**Figure 3:**
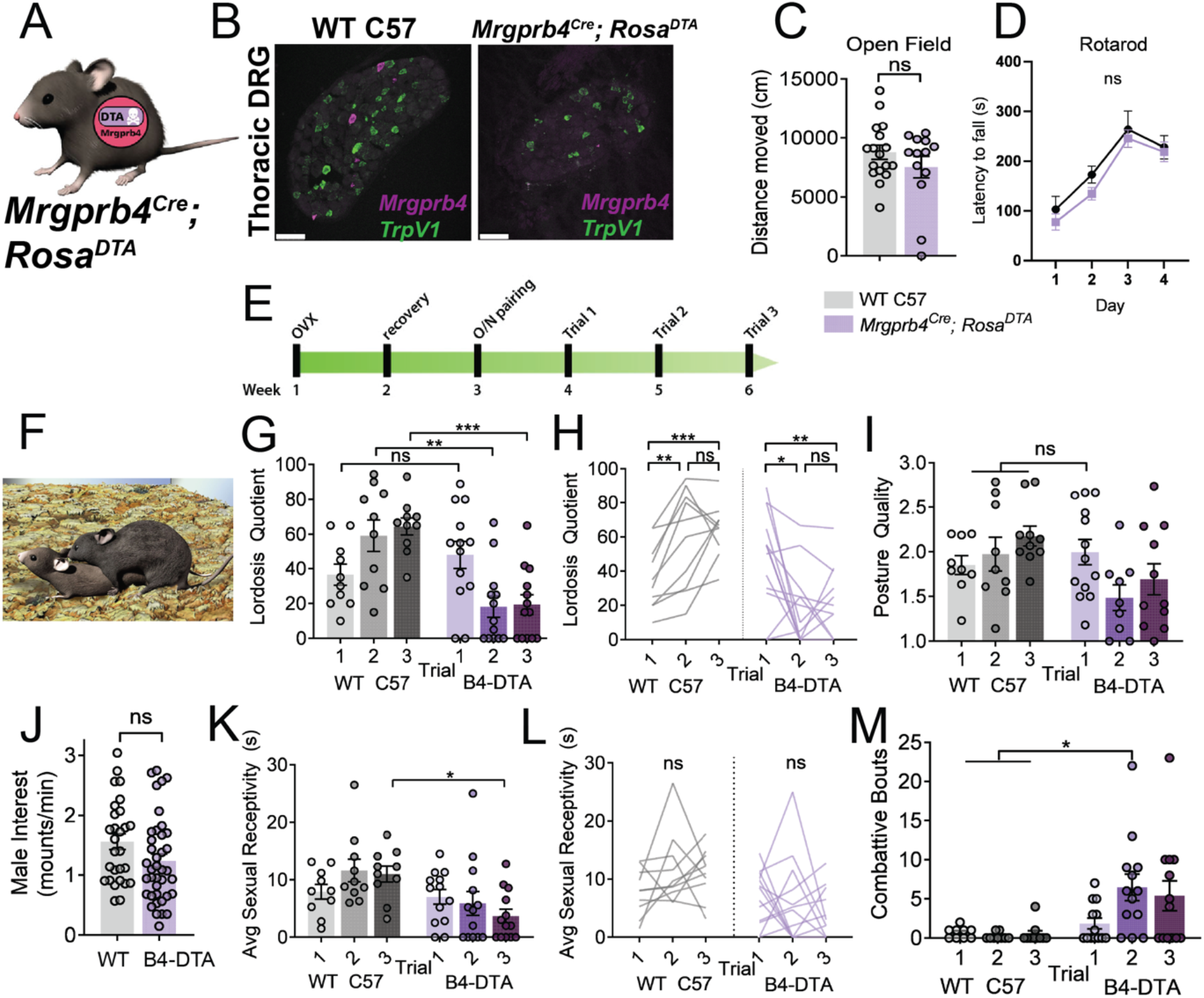
Mrgprb4-lineage neurons are required for female sexual receptivity. A) Graphic representing *Mrgprb4*^*Cre*^; *Rosa*^*DTA*^ mouseline, expressing DTA in a Cre-dependent manner to ablate Mrgprb4-lineage neurons. B) RNAscope *in situ* hybridization of DRGs from B) WT C57 control and C) *Mrgprb4*^*Cre*^; *Rosa*^*DTA*^ female, cell bodies expressing *Mrgprb4* (magenta) are absent upon DTA expression while *TrpV1* (green) cells remain intact. Scale bars = 75µm. C,D) *Mrgprb4*^*Cre*^; *Rosa*^*DTA*^ mice exhibit no motor deficits in open field (unpaired t-test) or rotarod assays (2-way ANOVA). E) Timeline for assessing sexual receptivity with lordosis quotient (LQ) assay: all females are given two weeks to recover after ovariectomy, at which point estradiol and progesterone are administered to put them in behavioral estrus for an overnight pairing with a male. Hormones replaced prior to each LQ trial. F) Assays conducted in the male home cage in the dark cycle. Graphic depicts a sexually receptive female lordosis posture G) Lordosis quotient scores for 3 sequential trials for WT C57 and *Mrgprb4*^*Cre*^; *Rosa*^*DTA*^ females. (*\*P<0*.*05, **P<0*.*01, ***P<0*.*001*, One-Way ANOVA). H) Changes in individual mice across the three trials. (*\*P<0*.*05, **P<0*.*01, ***P<0*.*001*, Repeated Measures ANOVA). I) Posture quality assessed on a scale from 1-3, where 1 is the minimum receptive posture and 3 is the most robust posture (details in methods). J) Male mounting frequency was no different between *Mrgprb4*^*Cre*^; *Rosa*^*DTA*^ females and controls. K) Average sexual receptivity, or duration maintained a receptive posture. (*\*P<0*.*05*, One way ANOVA). L) Changes in sexual receptivity in individual mice across trials (Repeated measures ANOVA). M) Total number of combative bouts observed for each trial. By the third trial, *Mrgprb4*^*Cre*^; *Rosa*^*DTA*^ females exhibited significantly more combative bouts than control females on any trial (\**P<0*.*05*, One way ANOVA).

To determine whether *Mrgprb4*^*Cre*^; *Rosa*^*DTA*^ females resisted mounts on subsequent pairings because they were incapable of producing a quality lordosis posture, we examined the quality of the posture in each of the three weeks. We found that their posture quality was normal on the first trial, suggesting that *Mrgprb4*^*Cre*^; *Rosa*^*DTA*^ females are indeed capable of producing a quality lordosis posture, but choosing not to display lordosis on subsequent trials (Fig 3i). Despite the reduced sexual receptivity of females, males demonstrated equal vigor in mounting in both *Mrgprb4*^*Cre*^; *Rosa*^*DTA*^ females and WT C57 controls (Fig 3j). Concomitant with the sharp decline in sexual receptivity in *Mrgprb4*^*Cre*^; *Rosa*^*DTA*^ females, these females now displayed combative behavior towards the male attempts to mount (Fig 3m). This phenotype is striking, as female mice primed for behavioral estrus are not naturally aggressive(Brock et al., 2011; Frye et al., 2014). Mrgprb4-lineage neurons are therefore not required for a motor output of lordotic dorsiflexion, but must encode a sensation that is necessary for the rewarding nature of sexual touch.

The “conspecific crawl” behavior, in which one female adopts a dorsiflexion posture as she crawls beneath a cage mate (thereby receiving social back contact) (Fig 4a), reminded us of postures we observed with optogenetic stimulation of Mrgprb4-lineage neurons (Fig 4b). The resemblance between natural and optogenetically evoked postures led us to ask if Mrgprb4-lineage neurons are required for female-to-female social behavior that involves back touch. We found that when *Mrgprb4*^*Cre*^; *Rosa*^*DTA*^ adult females and WT-C57 cage mates were observed for spontaneous social interaction, there was a specific deficit in social touch-induced dorsiflexion, or conspecific crawl (Fig 4e). The conspecifc crawl was a dorsiflexion response to any social touch to the back: allogrooming, other taps or contacts to the back, or sliding underneath a stationary cage mate (example graphic Fig 4d). Self-grooming, initiating social touch, allogrooming, and nestmaking behaviors were similar between groups (Fig 4f-j). Relatedly, *Mrgprb4*^*Cre*^; *Rosa*^*DTA*^ mice display normal social interest as tested in a social approach assay when presented with the choice of interacting with another mouse or an inanimate object (Fig 4h). Thus, Mrgprb4-lineage neurons are not necessary for typical social development but are necessary for evoking touch-dependent social postures.

**Figure 4:**
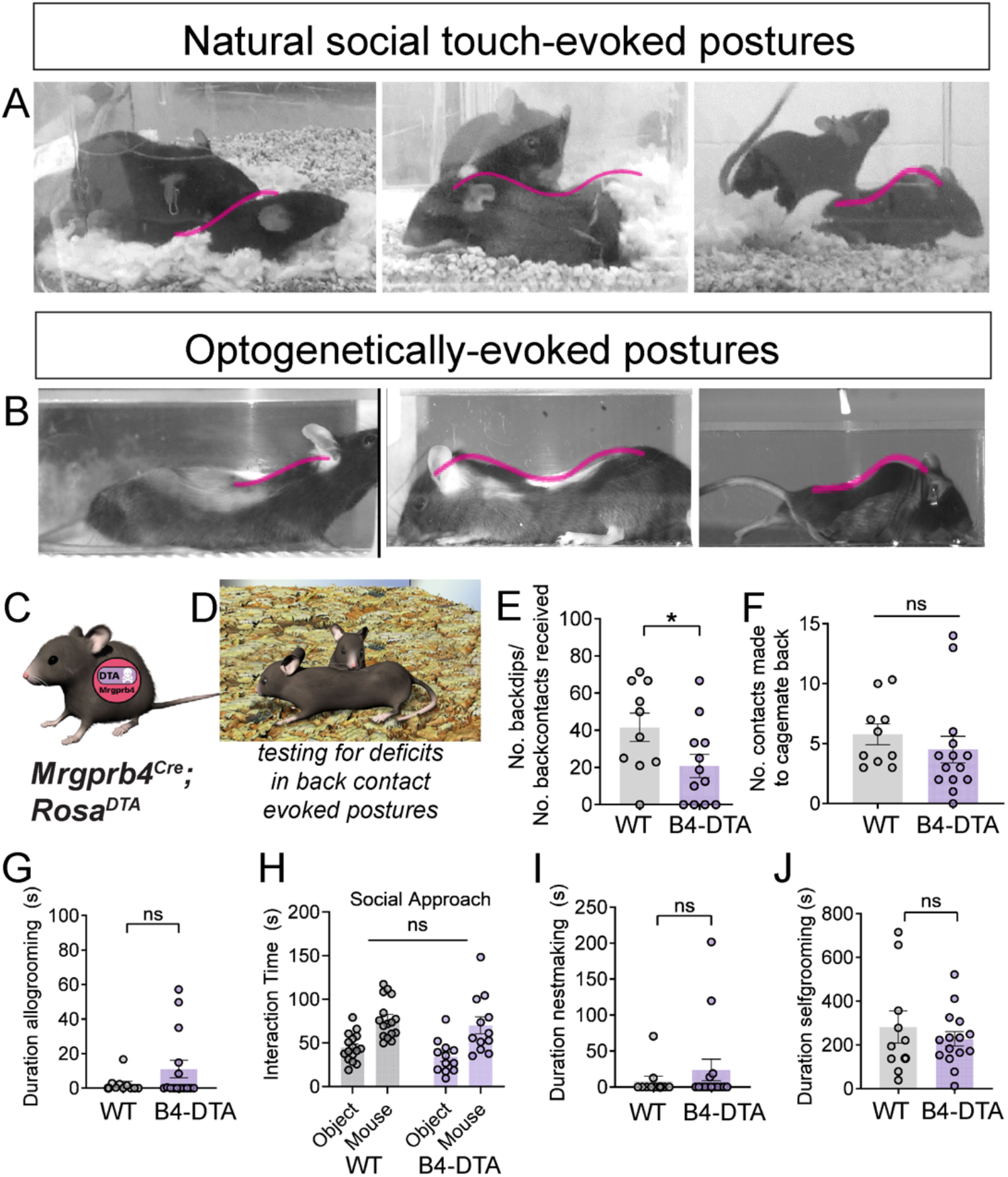
Mrgprb4-lineage neurons are required for female-to-female touch-evoked social postures but not general social interest. A) Still frames from behavioral videos showing the conspecific crawl posture: dorsiflexion in response to social touch. B) Still frames from highspeed videos showing the dorsiflexion induced by optogenetic activation of Mrgprb4-lineage neurons. These stills are eerily reminiscent of the postures induced by natural social touch. C) The *Mrgprb4*^*Cre*^; *Rosa*^*DTA*^ females were used in this experiment. D) Graphic depicting a typical conspecific crawl back dip posture E) Number of conspecific crawl back dip occurrences per mouse divided by the total number of backcontacts that mouse received. *Mrgprb4*^*Cre*^; *Rosa*^*DTA*^ females do fewer back dips than WT females (*\*P<0*.*05*, unpaired t-test). F) *Mrgprb4*^*Cre*^; *Rosa*^*DTA*^ females contact their cagemates’ backs just as frequently as WT females contact their cagemates’ backs in the 20 minute trial. G) *Mrgprb4*^*Cre*^; *Rosa*^*DTA*^ females allogroom their cagemates for similar duration as controls. H) *Mrgprb4*^*Cre*^; *Rosa*^*DTA*^ females demonstrate similar preference for a mouse over an inanimate object compared to controls. I) *Mrgprb4*^*Cre*^; *Rosa*^*DTA*^ females spend a similar duration nestmaking compared to controls. J) *Mrgprb4*^*Cre*^; *Rosa*^*DTA*^ females spend a similar duration self-grooming compared to controls.

### Chemogenetic activation of Mrgprb4-lineage neurons is not sufficient to promote social interaction or postures

Next, we sought to determine whether activation of the Mrgprb4-lineage neurons could increase the frequency of conspecific crawl postures in a natural social setting. We therefore used a Designer Receptors Exclusively Activated by Designer Drugs (DREADD)-based chemogenetic approach to activate Mrgprb4-lineage neurons while avoiding experimenter intervention during the social paradigm, which would be required for transdermal optogenetic activation. We generated *Mrgprb4*^*Cre*^; *Rosa*^*hM3dq*^ mice and either *Mrgprb4*^*Cre*^ *or Rosa*^*hM3dq*^ littermate controls and administered Clozapine N-oxide (CNO) or saline 30 minutes prior to reuniting the cage mates for the social interaction assay, as performed in Fig 3 (Supplemental Figure 4a-d). We observed that chemogenetic activation of Mrgprb4-lineage neurons did not promote increased social contact, an unsurprising result given that DTA-mediated ablation of Mrgprb4-lineage neurons did not impair social motivation or social contact (Supplemental Figure 4f,g). This further highlighted that Mrgprb4-lineage neurons, although recruited in touch-dependent social behaviors, do not appear to play a role in social motivation among same-sex conspecifics. Interestingly, however, there was no increase in social conspecific crawl postures (Supplemental Figure 4e). While optogenetic activation of Mrgprb4-lineage neurons is sufficient to induce a dorsiflexion posture in isolation, chemogenetic activation was not sufficient to promote the posture in social settings. It may be that focal activation on the back is required to generate the dorsiflexion, and body-wide chemogenetic activation is not a physiologically relevant sensory input for this social posture.

### Optogenetic-induced dorsiflexion posture is not modulated by ovarian hormones

Decades of work have established that rodent lordosis is tightly modulated by levels of estrogen and progesterone(Larsson et al., 1976; White and Uphouse, 2004; Yanase and Gorski, 1976). To determine whether the dorsiflexion caused by optogenetic activation of Mrgprb4-lineage neurons posture represents a posture of sexual receptivity, we tested whether the posture was modulated by ovarian hormones. First, we tracked the extent of dorsiflexion with the natural estrous cycle determined by daily vaginal lavage. We found that the dorsiflexion posture during 20s of optogenetic stimulation did not significantly vary across estrous states (Supplemental Figure 5a). Secondly, we determined whether experimental manipulation of ovarian hormones would impact the posture. We measured the optogenetic-induced postures in either vehicle or hormone-replaced ovariectomized females (to mimic a natural state of behavioral estrus). Consistent with the results for the natural estrous cycle, we found that the extent of the posture was not significantly different in ovariectomized compared to OVX + hormone replaced females (Supplemental Figure 5b). Thus, the dorsiflexion that results from the optogenetic activation of Mrgprb4-lineage neurons does not represent a state of sexual receptivity. Instead, the optogenetic activation of Mrgprb4-lineage neurons in the back skin may be triggering a local sensorimotor circuit that is recruited and modulated by hormonal conditions during natural sexual encounters.

**Figure 5:**
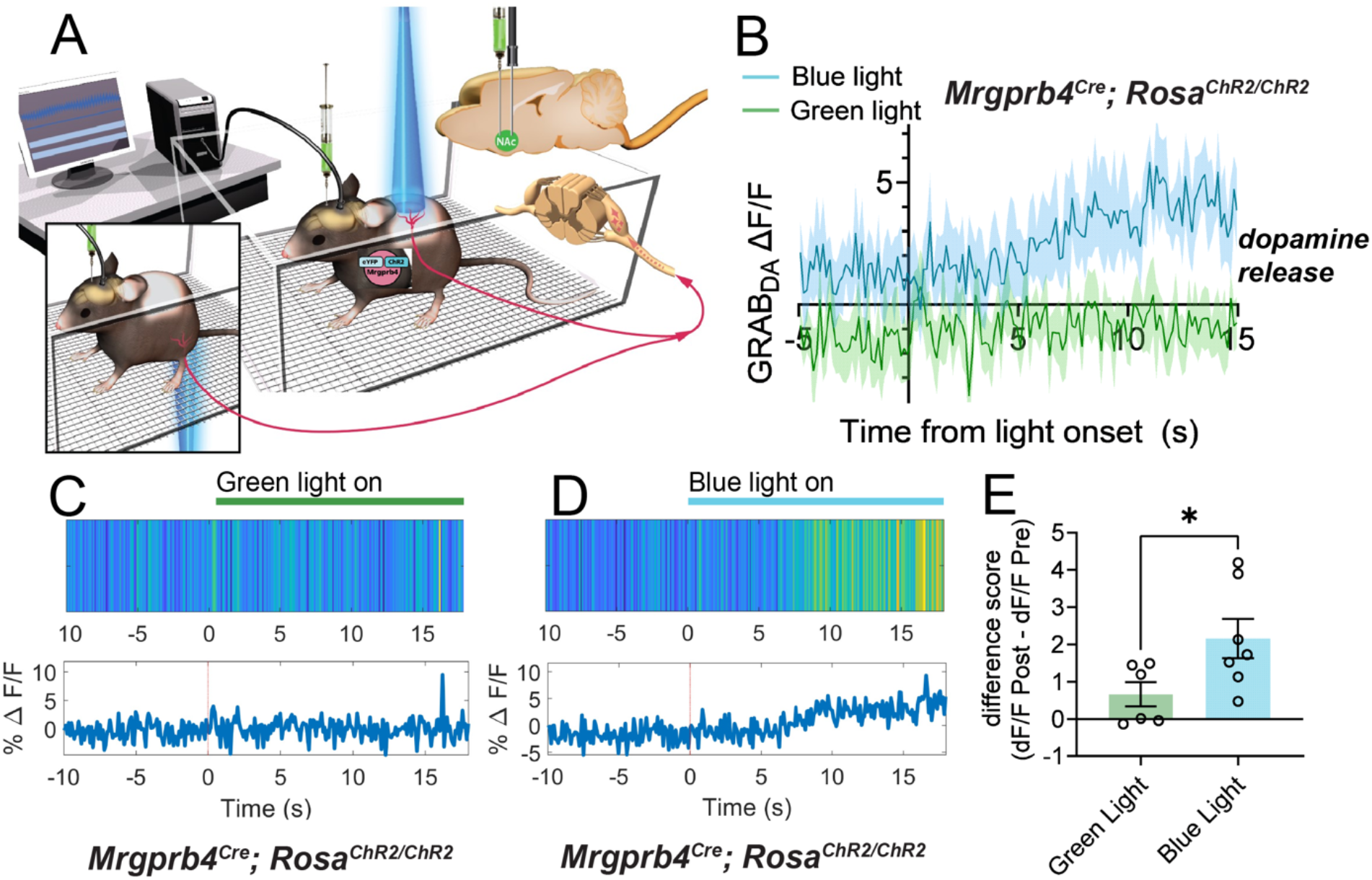
Activation of Mrgprb4-lineage neurons triggers dopamine release in the nucleus accumebns. A) Schematic depicting the experimental setup. Female *Mrgprb4*^*Cre*^; *Rosa*^*ChR2/ChR2*^ mice with shaved backs, which had been injected with GRAB_DA_ to the NAc two weeks prior to testing, were placed under plastic chambers on a mesh platform. Pulsed blue (stimulating) or green (control) laser light (35mW, 10Hz sin wave) was shined to either the back skin or skin surrounding the vagina while recording GRAB_DA_ signals. B) Average GRAB_DA_ delta F/F signals (N=6-7) for green and blue light. C) Representative trace from a mouse with green light shined to the skin surrounding the vagina, light stimulation begins T=0. D) Representative trace from the same mouse with blue light shined to the skin surrounding the vagina. E) Average deltaF/F pre stim (−5-0) subtracted from average deltaF/F post stim (10-15s) is significantly greater for blue light stimulation compared to the green light control, (\**P<0*.*05*, unpaired t-test).

### Optogenetic activation of Mrgprb4-lineage neurons is sufficient to cause dopamine release

We hypothesized that Mrgprb4-lineage neurons may directly transduce rewarding sensation during sexual encounters. Because tactile input from the male to the female during sexual behavior occurs to the back and anogenital region, we examined the skin in both regions for Mrgprb4+ nerve endings. Consistent with prior neuroanatomical studies (Liu et al., 2007), Mrgprb4+ terminals innervated all examined hairy skin, including the back, underbelly, and skin surrounding the genitalia, but were absent from the glabrous lining of the vaginal wall (Figure 1r; Supplemental Figure 6). We next coupled fiber photometry with transdermal optogenetic stimulation to test whether activation of Mrgprb4-lineage neurons – in either the back or anogenital skin – is sufficient to induce dopamine release in nucleus accumbens (NAc), a brain region for dopaminergic sexual reward(Becker et al., 2001; Beny-Shefer et al., 2017; Jenkins and Becker, 2003; Pfaus et al., 1995) (Fig 5a). To measure dopamine release we used stereotactic viral injections of the GRAB_DA_ dopamine sensor into the NAc. The GRAB_DA_ sensor is a GPCR-based approach where dopamine release and binding to its cognate receptor generates fluorescence with subcellular resolution and sub-second kinetics that we can detect with a photodetector (Sun et al., 2018). *Mrgprb4*^*Cre*^; *Rosa*^*ChR2/ChR2*^ females expressing the GRAB_DA_ dopamine sensor in NAc were given either 30s transdermal optogenetic stimulation to the back or anogenital region or the same treatment with non-stimulating green light while dopamine signals were recorded. When we analyzed the GRAB_DA_ signal upon optogenetic back stimulation of Mrgprb4-lineage neurons, we did not observe a significant increase in dopamine release (Supplemental Figure 7). We next turned to the Mrgprb4-lineage neurons innervating the skin surrounding the genitalia, which are expected to be stimulated during male intromission into the vagina during mounting. Intriguingly, blue light stimulation to skin surrounding the genitalia, but not green, was sufficient to raise GRAB_DA_ deltaF/F significantly from baseline levels (Fig 5b-e). This demonstrates that the Mrgprb4-lineage neurons in the anogenital skin are sufficient, in otherwise socially-isolated mice, to induce dopamine release.

### Mrgprb4-lineage neurons are required for dopamine release during sexual behavior

The Mrgprb4-lineage neurons are both necessary for the positive reinforcement of female sexual receptivity and sufficient to cause dopamine release in isolated females. We hypothesized that Mrgprb4-lineage neurons may be driving dopamine release during sexual behaviors. To determine whether Mrgprb4-lineage neurons are indeed required for sexual reward, we injected *Mrgprb4*^*Cre*^; *Rosa*^*DTA*^ females and littermate controls with the GRAB_DA_ dopamine sensor to observe any differences in dopamine activity during sexual behavior in the absence of Mrgprb4-lineage neurons (Fig 6a). We measured dopamine activity in the seconds surrounding mounts, anogenital sniffs, and other instances of back contacts to the female. We found that *Mrgprb4*^*Cre*^; *Rosa*^*DTA*^ females had significantly reduced dopamine signal immediately following mount initiation compared to littermate controls, observed in individual animals (Fig 6c,d), or averaged across animals (Fig 6b,e), suggesting the Mrgprb4-lineage neurons are indeed required for the sexual reward that occurs during a male mount. No significant difference in dopamine activity was detected for anogenital sniffs or back contacts (Supplemental Figure 8). Mrgprb4-lineage neurons may signal reward in these social touch scenarios, but dopamine release may be at levels too low to detect a difference, compared to the robust dopamine release in females upon male mounts. Together, these data reveal that Mrgprb4-lineage neurons communicate with the brain’s reward center during socially rewarding sexual behavior.

**Figure 6:**
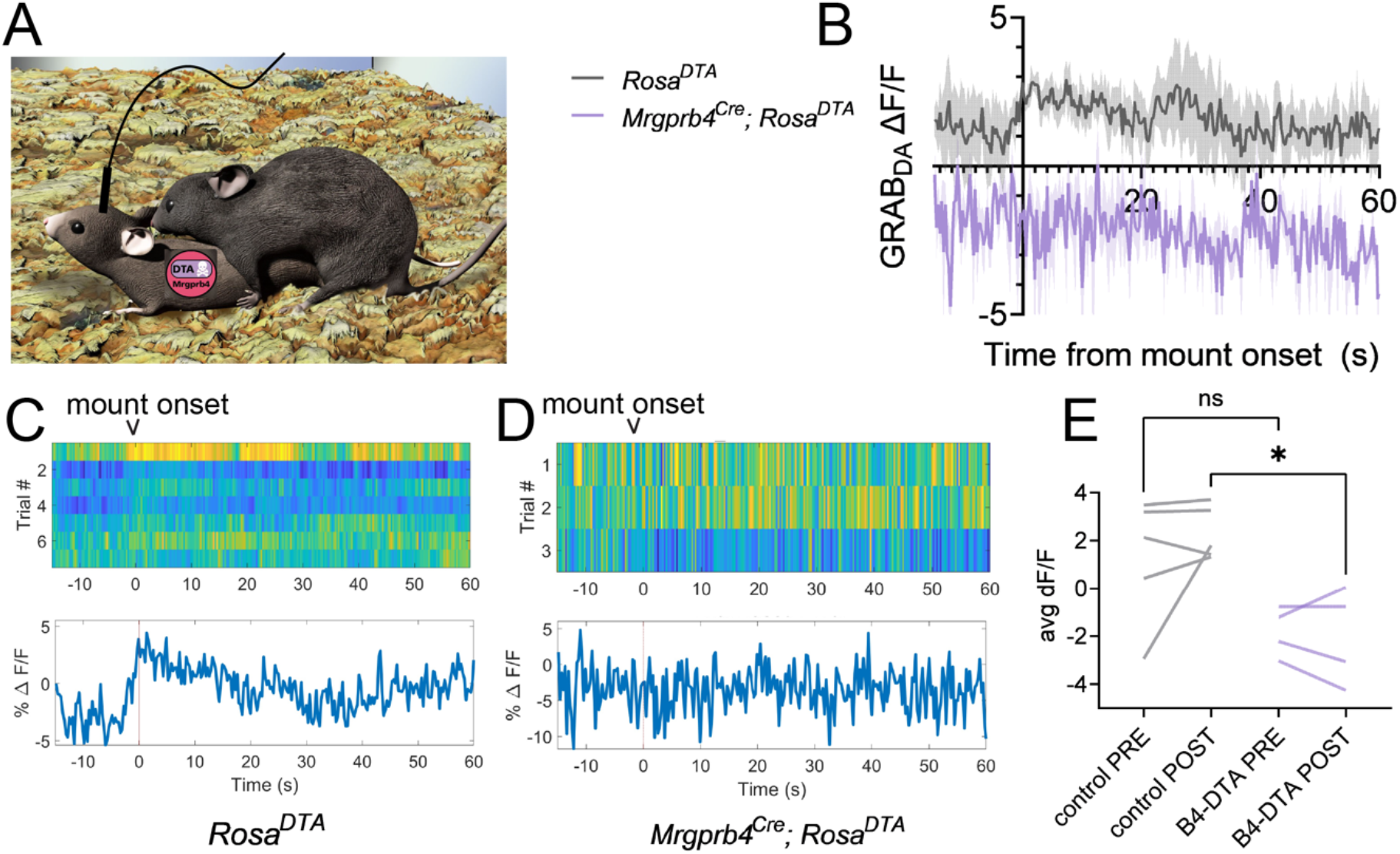
Mrgprb4-lineage neurons are required for dopamine release in the nucleus accumbens during sexual mounts. A) Graphic depicting experimental setup. Female *Mrgprb4*^*Cre*^; *Rosa*^*DTA*^ mice or littermate *Rosa*^*DTA*^ controls, which had been injected with GRAB_DA_ to the NAc over two weeks prior to testing were paired with males for a mating assay. Females were ovariectomized and hormone primed to be in a state of behavioral estrus for testing. GRAB_DA_ signals were recorded for the entire pairing and analyzed surrounding three events: mount, anogenital sniff to female, and back contact to female. B) Average deltaF/F traces surrounding mount onset (T=0) for *Mrgprb4*^*Cre*^; *Rosa*^*DTA*^ females (purple) or littermate *Rosa*^*DTA*^ controls (gray) (N=4-5). C) Representative average GRAB_DA_ signal across seven mounts in one pairing for a *Rosa*^*DTA*^ control, mount onset at T=0. D) Representative average GRAB_DA_ signal across three mounts in one pairing for a *Mrgprb4*^*Cre*^; *Rosa*^*DTA*^ female. E) The average GRAB_DA_ signal following mount onset is reduced in *Mrgprb4*^*Cre*^; *Rosa*^*DTA*^ females compared to *Rosa*^*DTA*^ controls (\**P<0*.*05*, One way ANOVA).

## Discussion

Although affective social touch begins at the skin’s surface, the molecular identity of sensory neurons in the skin that detect socially relevant signals and pass them to the central nervous system has remained unknown. Moreover, because touch itself is highly heterogeneous (i.e., discriminative touch to detect texture with our fingertips versus the affiliative touch during a hug from a friend), sensory perception is likely generated by different sets of neurons to provide specificity (Handler and Ginty, 2021; Li et al., 2011; Maksimovic et al., 2014; Neubarth et al., 2020; Rodgers et al., 2021; Seal et al., 2009b; Severson et al., 2017; Sharma et al., 2020). Armed with this information, where does one begin the search for touch neurons underlying social reward, including sexual receptivity? Within the deep ocean of DRG neuron types, one class, termed C-low threshold mechanoreceptors (C-LTMRs), or C-tactile afferents in humans, are implicated in detecting gentle strokes across the skin’s surface(Ackerley et al., 2014). Although a limited number of papers in the mouse identify molecular populations of C-LTMRs, including their neuroanatomy and roles in somatosensation during baseline and chronic pain states (Bohic et al., 2020; Francois et al., 2015; Kuehn et al., 2019; Li and Ginty, 2014; Li et al., 2011; Lou et al., 2013; Seal et al., 2009a; Urien et al., 2017), the role of C-LTMRs in promoting social behaviors remains obscure.

Here, we demonstrate that Mrgprb4-lineage neurons are indeed critical for specific social behaviors and for signaling social reward to the brain. Focal activation of Mrgprb4-lineage neurons yields a striking dorsiflexion posture resembling mammalian lordosis, representing the first acute behavioral response to optogenetic activation of social touch neurons. Mrgprb4-lineage neurons are required for two touch-dependent social postures: sexual behavior and crawling under same-sex conspecifics. However, their role in female sexual behavior is not simply in the local lordotic reflex; rather, the neurons convey an affective sensation that reinforces sexual receptivity, and without them, male advances during sexually receptive hormonal states become aversive. This finding suggests that Mrgprb4-lineage neurons contribute to the perceived valence of sexual encounters in females by encoding the rewarding aspect of male sexual touch. Lastly, we use fiber photometry to functionally link Mrgprb4-lineage neurons in the skin to reward circuitry in the brain. Transdermal optogenetic activation of Mrgprb4-lineage neurons in the skin surrounding female genitalia is sufficient to induce dopamine release in the NAc: this is the first report of molecularly defined somatosensory neurons triggering activation of a brain reward center. Finally, we link this simulation of rewarding touch to natural tactile reward by demonstrating that Mrgprb4-lineage neurons are required for this same dopamine release during male mounts.

One question that emerges from our genetic targeting strategy is whether the effects that we have revealed could be driven solely by the Mrgprb4+ adult population of neurons, or whether the effects are attributed to the broader population of sensory neurons that share developmental history of expressing Mrgprb4. Moreover, is there an unidentified gene or molecule expressed in Mrgprb4-lineage neurons, that might be important in initiating the pleasurable touch circuit in the skin? We believe the studies here open the field to explore these and other exciting new possibilities, particularly the importance of considering lineage expression of a gene a property that can define a functional class of neurons.

In summary, the work described here has revealed the sufficiency of peripheral inputs to regulate social reward independent of context and other sensory cues. Since ventral tegmental dopaminergic neurons that project to the NAc are themselves functionally heterogeneous (Lammel et al., 2011; Morales and Margolis, 2017), it is possible that some of these neurons might be tuned for encoding rewarding touch. It will also be interesting to determine if Mrgprb4-lineage neurons are involved in other behaviors that integrate social touch, such as maternal care. Another outstanding question in touch biology is whether defined molecular classes of touch neurons are tuned for encoding unique behaviors or whether a combinatorial code of touch neurons is activated in the skin with specificity driven by central circuits. The approaches outlined here genetically targeting touch neurons during behavior and imaging could be leveraged to determine the functional roles of other molecularly defined touch neurons. We believe this study draws new attention to the importance of elucidating skin-brain circuits, analogous to the importance of gut-brain circuits. Moreover, this work points towards the therapeutic potential of peripheral manipulations for enhancing intact or impaired social reward systems, including sexual receptivity, or simulating social reward during periods of isolation.

---

**Supplemental Figure 1.**
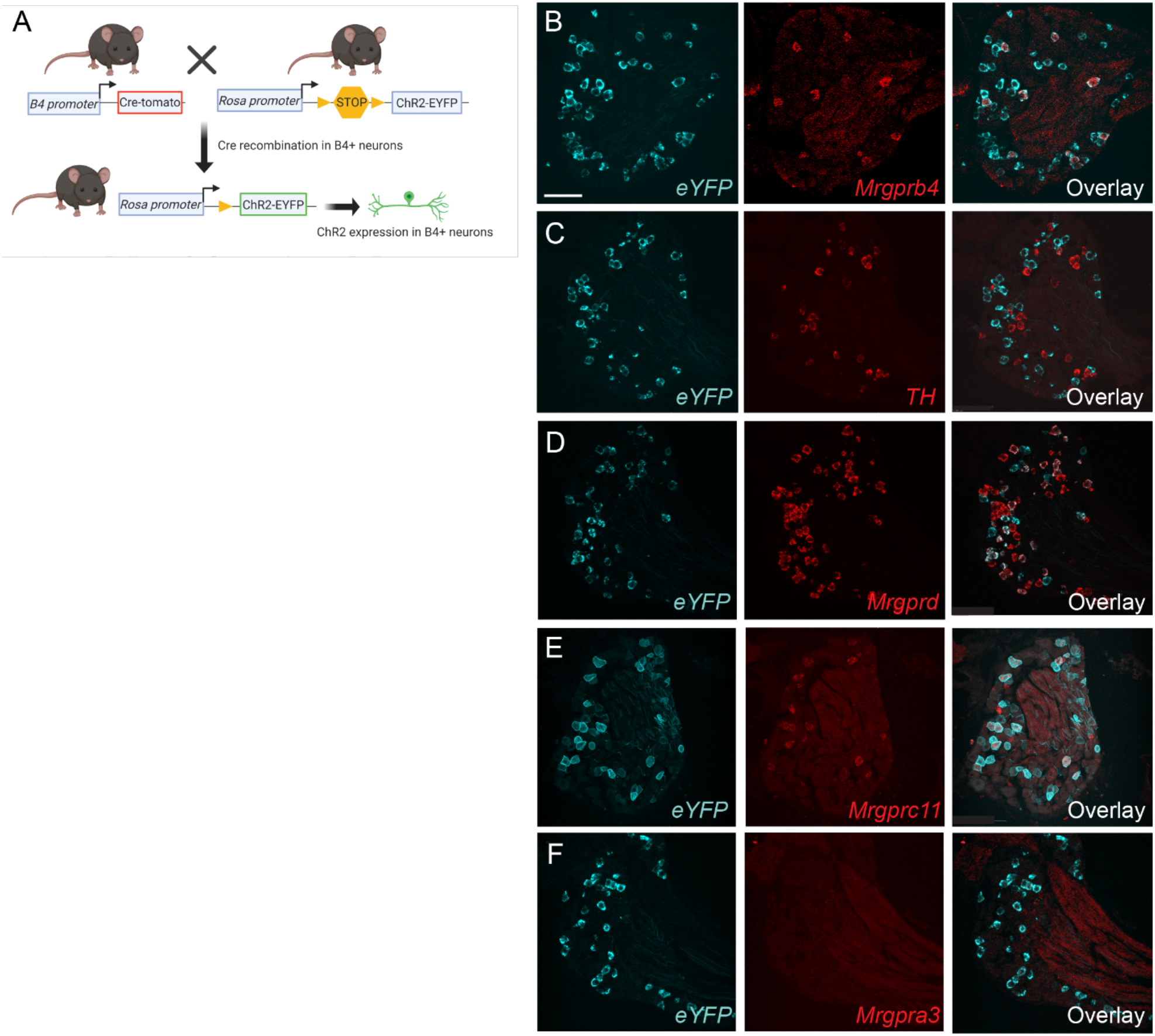
Characterizing expression of ChR2-eYFP in Mrgprb4-lineage neurons. A) Schematic showing breeding strategy to express ChR2-eYFP in Mrgprb4-lineage neurons. B-F) Double RNA *in situ* hybridization with RNAscope probes that detect eYFP (cyan) or another marker gene (red). Quantification and description of results in Figure 1d,e. Scale bar represents 100 µm.

**Supplemental Figure 2.**
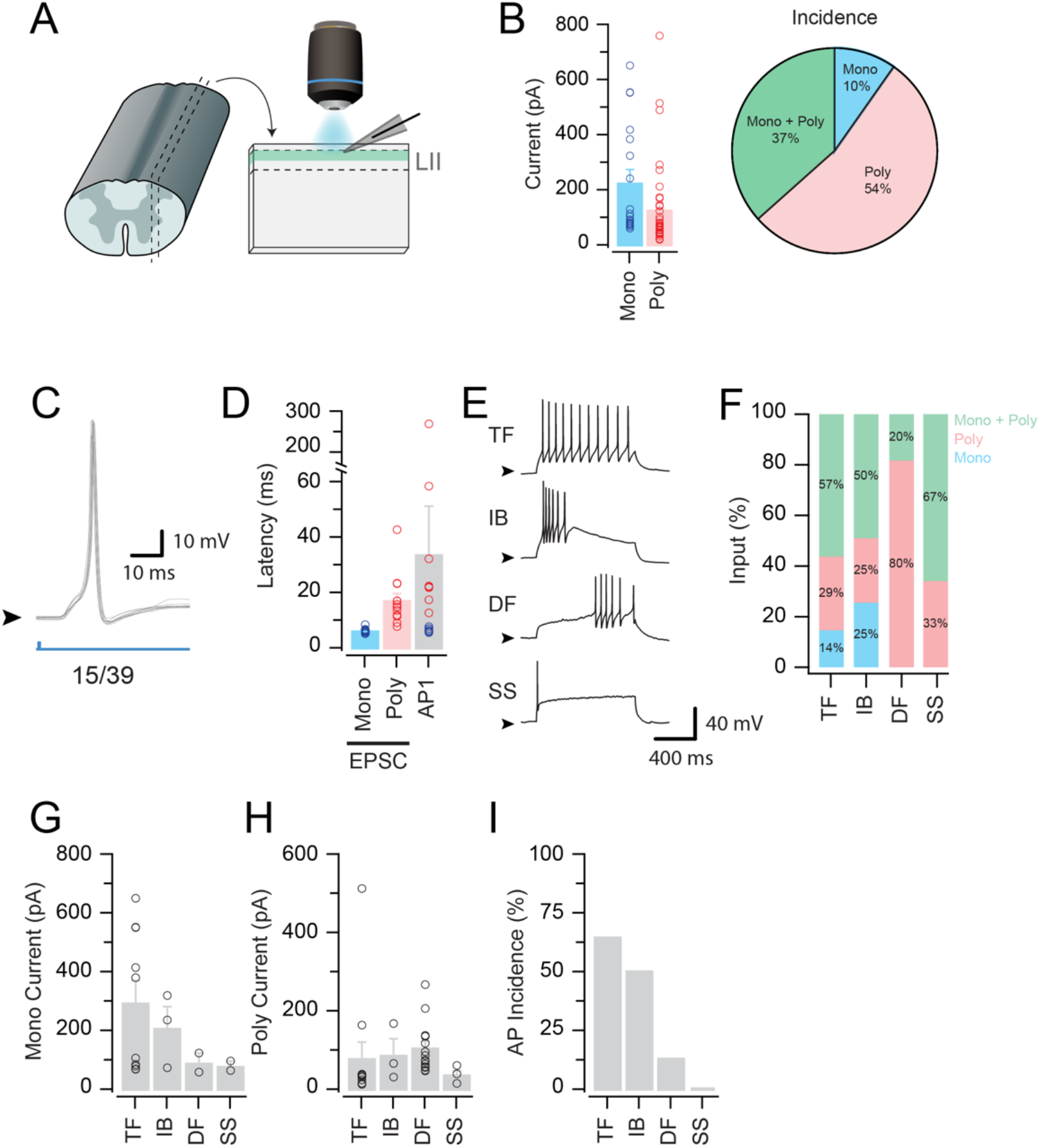
Characterizing physiological properties of dorsal horn spinal cord neurons that receive synaptic input from Mrgprb4-lineage neurons. A) Schematic of the spinal cord slice preparation to activate Mrgprb4+ terminals with blue light in the superficial dorsal horn and record from synaptically connected neurons. B) Left shows grouped data of mono and polysynaptic EPSC amplitudes. Right shows incidence of a monosynaptic only, polysynaptic only, or mono + polysynaptic input. C) Optically evoked action potential (AP). 15/39 neurons tested fired AP’s following 1ms photostimulation. D) Latency of oEPSCs (mono and polysynaptic) in these neurons, compared to the latency of the first evoked AP. Of these 15 neurons, 7 display characteristics of AP’s evoked by direct Mrgprb4 afferent input, 8 display characteristics of AP’s evoked by indirect (polysynaptic) Mrgprb4 inputs. E,F) In a subset of recordings AP discharge was characterized in post-synaptic neurons (n = 36). 2/14 Tonic firing (TF) neurons received mono input only, 4/14 poly only, and 8/14 both. 0/15 Delayed firing (DF) neurons received mono input only, 12/15 received poly only, and 3/15 both. 1/4 Initial bursting (IB) neurons received mono input only, 1/4 received poly only, and 2/4 both. 0/3 Single spikers (SS) received mono input only, 1/3 received poly only, and 2/3 both. G) TF neurons receive the strongest monosynaptic inputs, followed by IB’s. H) Polysynaptic currents are similar across all types. I) AP incidence is highest in TF neurons.

**Supplemental Figure 3.**
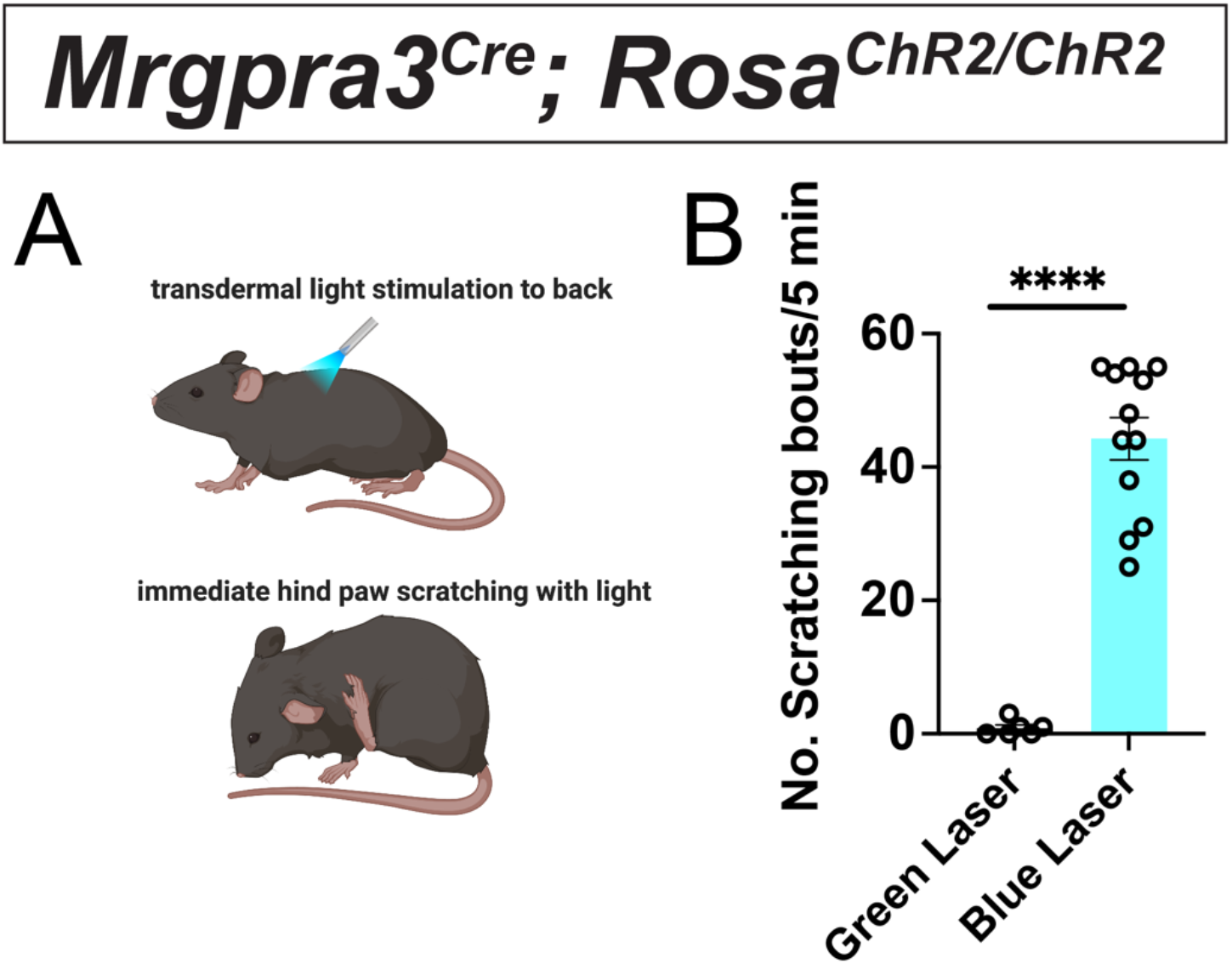
Transdermal optogenetic activation of Mrgpra3+ neurons elicits stereotyped scratching behaviors. A) Schematic showing experimental setup with blue light applied to the shaved back skin, which elicits nearly immediate scratching bouts, and not the back lowering dorsiflexion phenotype observed when the same experiments are performed with Mrgprb4-lineage neurons. B) Quantification of scratching bouts to 5 minutes of blue or green laser light (negative control). Unpaired student’s t-test, **** *P* ≤ 0.0001. Individual circles represent a single mouse.

**Supplemental Figure 4.**
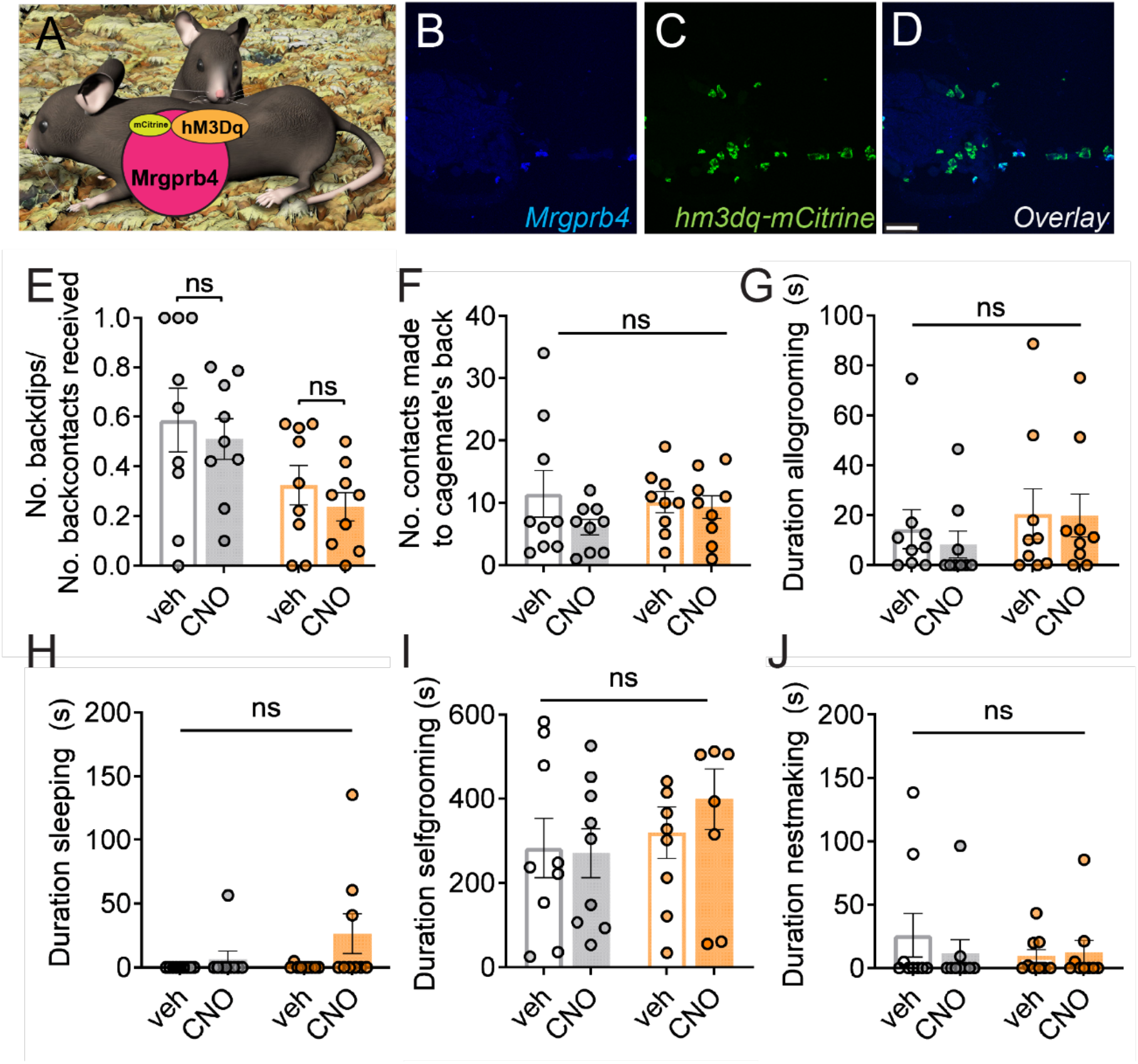
Body-wide chemogenetic activation of Mrgprb4-lineage neurons does not alter social behaviors. A) Schematic showing chemogenetic strategy to activate Mrgprb4-lineage neurons with DREADD-mediated activation while assessing social behaviors. B-D) Confirmation of targeting strategy to insert the DREADD receptor hM3Dq-mCitrine into Mrgprb4-lineage neurons. Data visualized with RNAscope *in situ* hybridization. Scale bar represents 75 µm. E-J) No statistical difference in any assay measuring social (E-G) or other (H-J) behaviors. All data plotted as mean +/-SEM, with one-way ANOVA followed by Tukey’s post-hoc comparisons.

**Supplemental Figure 5.**
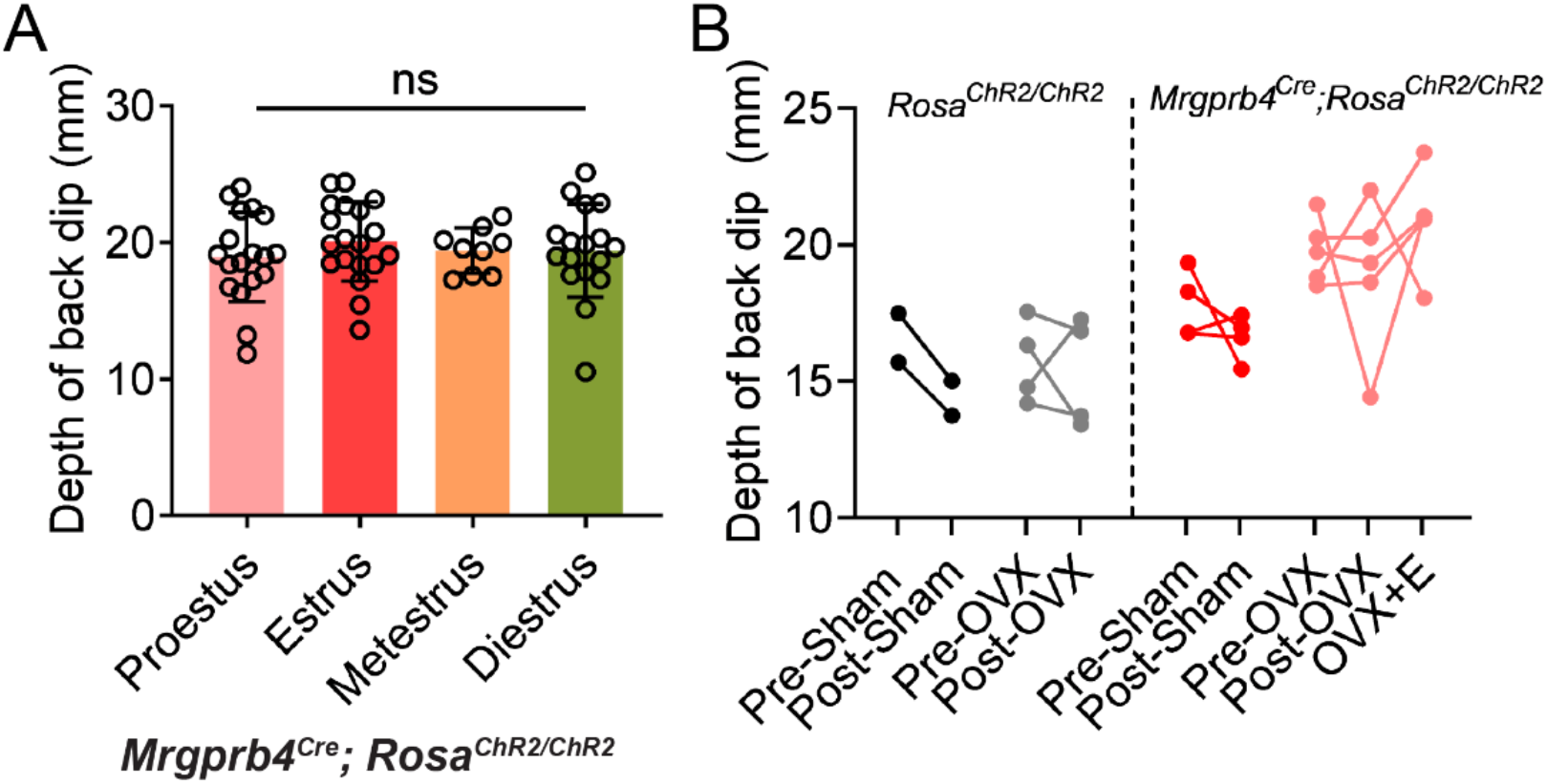
Lordosis-like dorsiflexion posture observed in *Mrgprb4*^*Cre*^; *Rosa*^*ChR2/ChR2*^ female mice is independent of estrous state. A) No differences in the depth of the optogenetically induced back dip across stages of the estrous cycle. Circles represent individual mice, data plotted as mean +/- SEM. B) Neither ovariectomizing (OVX) females, nor replacing estrogen and progesterone exogenously to mimic behavioral estrus, significantly modulates the optogenetically induced back dip. Dots represent individual mice. All data analyzed by one way ANOVA followed by Tukey’s post-hoc comparisons.

**Supplemental Figure 6.**
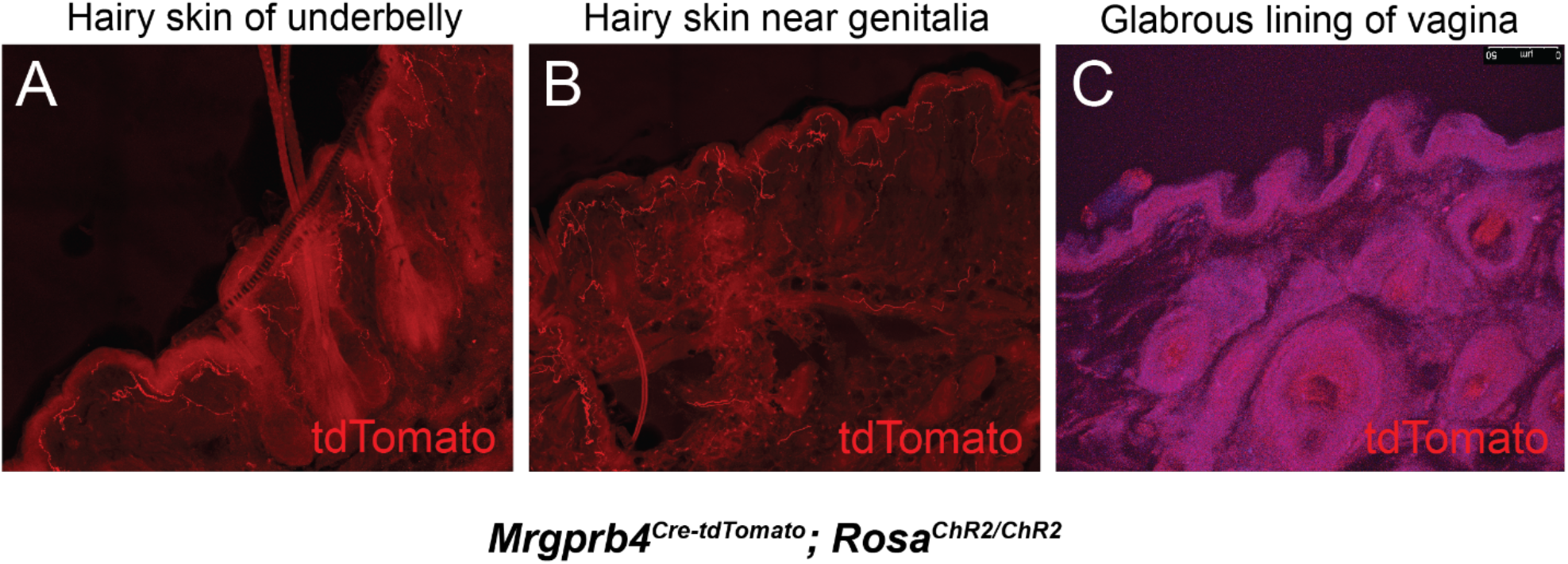
Nerve terminal endings of Mrgprb4+ neurons in skin of the anogenital regions with immunostaining of tdTomato. A,B) Nerve terminal endings in red are seen in the underbelly skin as well as the hairy skin surrounding the female genitalia. C) Nerve terminals for Mrgprb4+ touch neurons are not seen in the vaginal wall, which is glabrous, non-hairy skin.

**Supplemental Figure 7.**
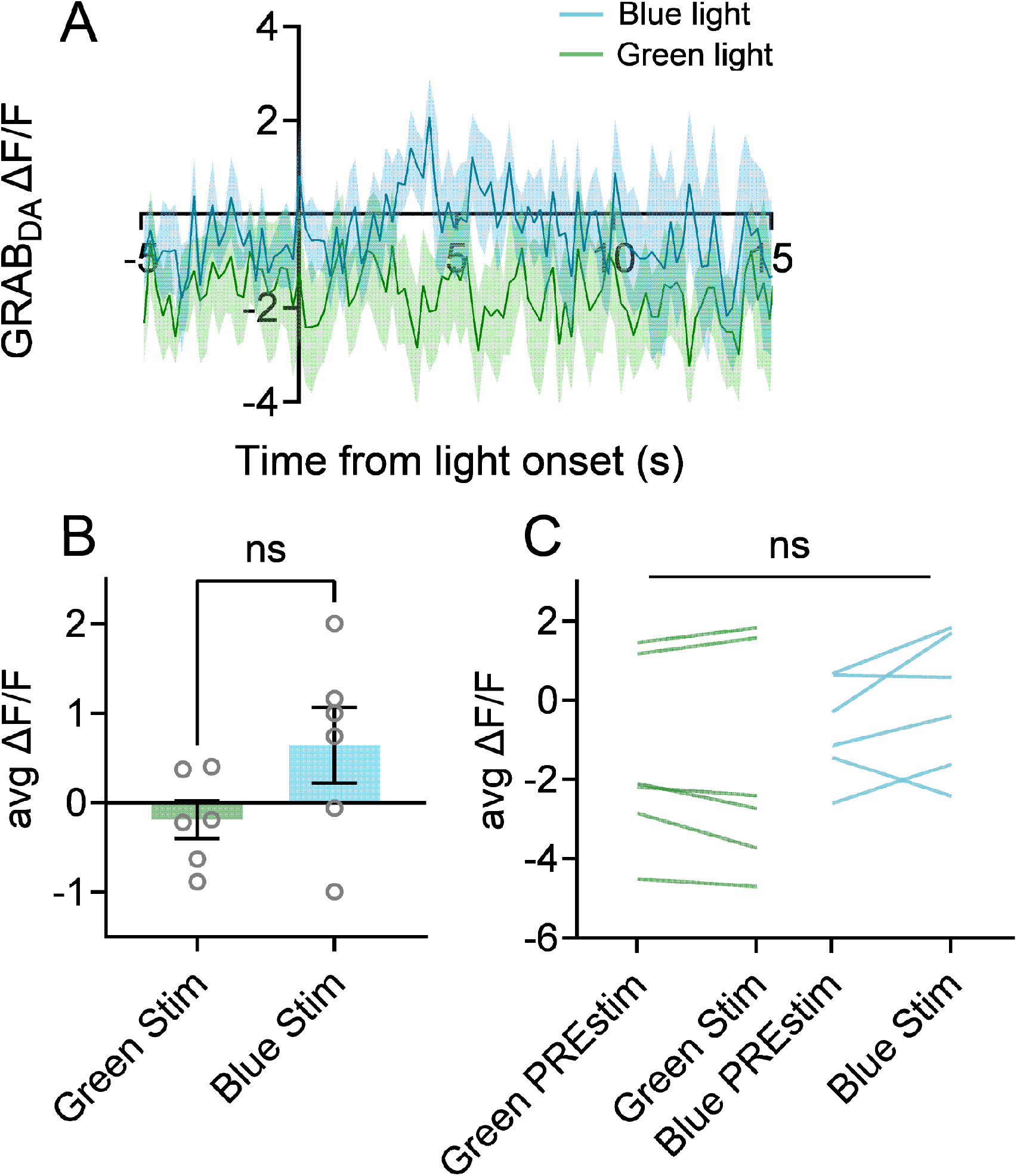
Optogenetic activation of Mrgprb4-lineage neurons in the back skin does not lead to detectable dopamine release. A) Average GRAB_DA_ delta F/F signals (N=6-7) for green and blue light. B) Average deltaF/F pre stim (−5-0) subtracted from average deltaF/F post stim (10-15s) is not statistically significantly different for blue light stimulation compared to the green light control, unpaired t-test. C) Average deltaF/F pre and post stim are not statistically different between blue and green light control stimulation, one-way ANOVA.

**Supplemental Figure 8.**
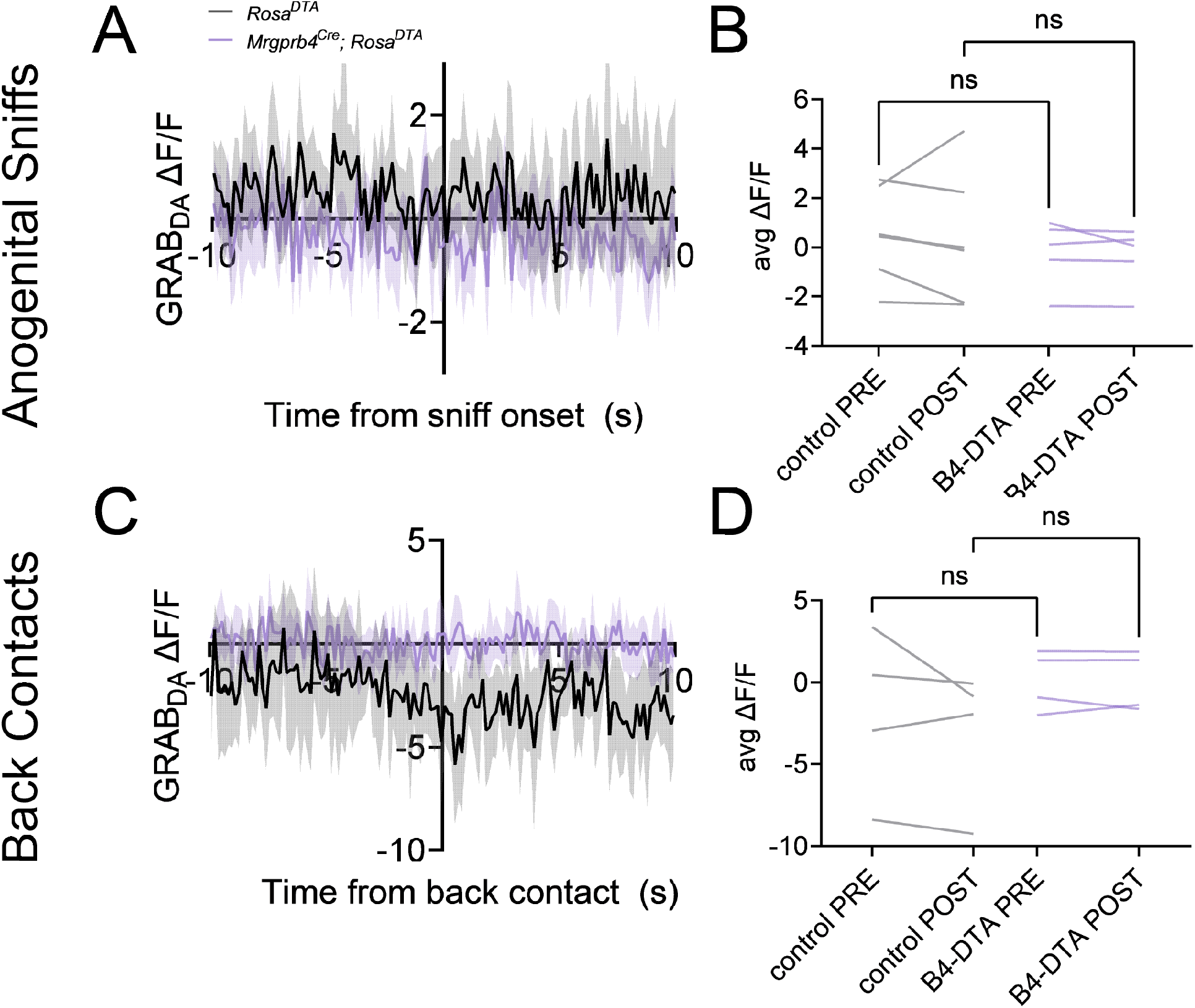
No changes in dopamine release in females observed in control or Mrgprb4-neuron ablated mice to anogenital sniffs or back contacts. A,C) Average deltaF/F traces (N=4-6) in the 20s surrounding (A) anogenital sniffs or (C) back contacts received from the male during the sexual encounter. B,D) Average deltaF/F pre (−5-0s) and post (0-10s) anogenital sniff (B) or back contact (D) are not statistically different between B4-DTA and control female mice, one-way ANOVA.

## Methods

### Mice and Behavioral Testing

All testing was performed in compliance with the NIH Guide for the Care and Use of Laboratory Animals and were approved by the Institutional Animal Care and Use Committee of the University of Pennsylvania and Columbia University. Mice born into our colony on a C57bl/6J background were maintained in conventional housing with food and water available ad libitum when not being tested, on a 12 hour light cycle beginning at 7:00, and all testing unless otherwise noted occurred during the light cycle in a room directly adjacent to the housing room in the animal facility. Mouse lines used in this study are located at Jackson Laboratories: *C57BL/6J* (Stock No: 000664), *Mrgprb4*^*Cre*^ (Stock No: 021077), *Mrgprd*^*Cre-ERT2*^ (Stock No: 031286), *Rosa*^*ChR2-eYFP*^ (Stock No: 024109), *Rosa*^*DTA*^ (Stock No: 009669), *Rosa*^*Gq-DREADD*^ (Stock No: 026220). The *MrgprA3*^*Cre*^ line was generously provided by Dr. Xinzhong Dong at Johns Hopkins University School of Medicine.

### RNAscope in situ hybridization

Brains were harvested immediately following transcardial perfusion and post-fixed in 4% PFA in PBS at 4°C for 24h. Brains were subsequently submerged in 30% sucrose in PBS at 4°C for 18h or until sunk, flash frozen in cryomold on dry ice in OCT, sectioned at 12µm directly onto Superfrost Plus Slides and stored in foil-wrapped slide box at −80°C until beginning Fixed Frozen RNAscope protocol (ACD).

Entire spinal columns were harvested immediately following transcardial perfusion and post-fixed in 4% PFA in PBS at 4°C for 24h. Either spinal cord or DRG were subsequently dissected and submerged in 30% sucrose overnight or until sunk. Tissue was flash frozen in cryomold on dry ice in OCT, sectioned at 12µm directly onto Superfrost Plus Slides, and stored in foil-wrapped slide box at −80°C until beginning Fresh Frozen RNAscope protocol (ACD). Note the fixation step was skipped in ACD’s Fresh Frozen protocol for these tissues.

### Immunohistochemistry

Brains, spinal cord, and DRG were collected and prepared in the same way as *in situ* hybridization. 30µm cryosections were cut directly onto Superfrost Plus Slides. Slides were frozen overnight in a slide box at −80°C. Slides were washed 3x 10min in PBS; 30 minutes in PBST; 1 hour in PBST with 5% normal donkey serum. Primary antibody, 1:1000 in PBST with 5% normal donkey serum, was applied to slides in a humidified chamber overnight at room temperature. Secondary antibody, 1:400 in PBST with 5% normal donkey serum, was applied to slides in a humidified chamber 1-2 hours at room temperature. Slides were washed 3x 10 minutes in PBS before applying mounting media and coverslip.

### Transdermal optogenetic activation of Mrgprb4-lineage neurons

Transdermal optogenetic stimulation of sensory neurons was performed as we previously described^14^. In brief, 8-16 week old female mice were habituated to mesh platform under a plastic chamber for 1 hour on each of two days prior to behavioral testing. Experimenter was present with lights, camera, and laser running during habituation to mimic entire sensory experience of test day. On the day of testing, the mice were habituated to the chambers for an additional 20 minutes before stimulation. 35mW blue laser light pulsed at 10 Hz sin wave was shined through the ceiling of the chamber to the shaved backs of mice for 20 seconds during high-speed video captured at 750fps.

### Tamoxifen injection for Mrgprd^Cre-ERT2^; Rosa^ChR2/ChR2^ mice

To induce expression of Cre recombinase in *Mrgprd*^*Cre-ERT2*^; *Rosa*^*ChR2/ChR2*^ mice, we intraperitoneally injected 0.5mg tamoxifen in 100µl sunflower seed oil in mice aged P10 or older. We repeated this injection daily for three days, so each mouse received three 0.5mg doses. Behavioral testing began at least two weeks after the third injection to allow for Cre expression.

### Quantification of the back dip

Optogenetic-induced dorsiflexion posture was calculated as the maximum back dip from the ceiling of the behavioral chamber for the duration of the 20s stimulation. High speed videos were analyzed in ImageJ to track the lowest point of the back throughout the 20s stimulation. The lowest y value was subtracted from the y value of the top of the chamber, and the value was converted from pixels to millimeters by determining the pixel height of the 4.5cm chamber for each video. It was decided that this was the most quantifiable way to characterize the dip. While duration or frequency of response require a subjective determination of when a back dip starts and stops – and therefore can be more challenging to compare to control animals – the back dip depth is the most objective measure because it allows us to compare between natural movement of the spine in controls, small back dips, and most drastic back dips. This measure encapsulates the full variability of response. We plotted as the percent of animals that concaved their back beyond 17mm, which was chosen as a threshold based on the spread of the data to represent a dipping of the back beyond typical movements.

### Conditioned Place Preference

8-14 week old *Mrgprb4*^*Cre*^; *Rosa*^*ChR2/ChR2*^ females or *Mrgprd*^*CRE-ERT2*^; *Rosa*^*ChR2/ChR2*^ and Cre-negative *Rosa*^*ChR2/ChR2*^ female littermates underwent a 9 day conditioned place preference paradigm. The apparatus consisted of two chambers, one paired with almond extract and a textured floor, the other coconut extract and a smooth floor. Additionally, one chamber wall had stripes, while the other had polka dots. These olfactory and visual stimuli were present throughout the 9 day paradigm to aid in the mouse’s encoding of different chambers. Days 1-3 were habituation: each mouse was allowed 20 minutes to explore the two chambered apparatus with no optogenetic stimulation. Days 4-8 were training: each mouse was allowed 20 minutes to freely move about the two-chambered apparatus, now receiving laser light stimulation to the back. In one chamber the mouse received 10Hz pulsed sin wave 35mW blue laser light to the shaved back, and the other chamber non-stimulating green light of the same parameters. Experimenter held the laser lights ∼1cm from the back skin for the duration that the mouse was in the chamber. The lasers were held with an extendable alligator clip to avoid casting body shadow on the chambers. Day 9 was test day: each mouse was allowed 20 minutes to freely move about the two-chambered apparatus in the absence of any laser light stimulation. Baseline preference was calculated for each mouse by averaging the duration spent in each chamber across the three habituation days. The conditioned preference was calculated for each mouse as the duration spent in each chamber on the test day. Percent change was calculated as percent time in blue light chamber after training – percent time in blue light chamber before training.

### Determination of natural estrous state

Natural estrous cycle was determined by vaginal lavage. Immediately following behavioral testing, the vagina was flushed with 20µl ultrapure water which was then pipetted onto a Superfrost plus slide for examination under dissecting scope. Mice were tested mid-morning each day to ensure the most accurate tracking. Lavage samples were assessed as described (Cora et al., 2015), and estrous state was recorded throughout the week to ensure normal cycling. Data from mice with lavage samples that could not be fit into a typical four or five day cycling pattern were excluded.

### Ovariectomy Surgery

Ovariectomies were performed on 8 week old female mice under 1.5-2% isoflurane using proper sterile technique. 5mg/kg oral or intraperitoneal meloxicam and 2mg/kg subcutaneous bupivacaine (at incision sites) were administered before surgery. A 0.5cm incision is made 1cm lateral to spinal cord, at the point where ribcage ends. An equivalent incision was made through the muscle wall. The white fat pad was exposed to identify the ovary, which was cauterized with a hemostat and scraped off with a scalpel. The fat pad was reinserted and 1-2 sutures closed the muscle wall. 2-4 sutures closed the skin. Meloxicam was administered 24 hours after surgery and mice were allowed two weeks to recover before behavioral testing.

### Lordosis Quotient Assay

10-14 week old ovariectomized females underwent two overnight pairings with stud males two weeks following surgery. To mimic behavioral estrus state at the time of pairing, females were subcutaneously injected with 0.5 µg estradiol benzoate in sunflower seed oil both 52 and 28 hours before pairing, and with 800 µg progesterone in sesame oil 4 hours before pairing. The females received the same hormone treatment prior to the lordosis quotient assay. Lordosis quotient assay was conducted in male home cage, 1-2 hours into the dark cycle. Video of the assay was recorded for 20 minutes or 20 attempted male mounts, whichever came first. Lordosis quotient was calculated as the number of female receptive responses divided by the number of attempted male mounts. A receptive response for the female was scored as all four limbs securely on the cage floor with no combative or escape behaviors. If the first male did not mount within 10 minutes, the female was moved to another male’s cage. For any given trial, up to three males may have been used. Receptive postures were scored for quality on a scale from 1-3 as follows: 1: limbs on the ground with no attempts to escape, neither dorsiflexion nor upturned nose. 2: Some dorsiflexion, no upturned nose. 3: Robust dorsiflexion and/or upturned nose. Posture scores were averaged within each trial.

### Interfemale Social Behavior Assay

The interfemale social behavior assay was adapted from allogrooming assays(Burkett et al., 2016; Lu et al., 2018) and conducted in the same way for both diphtheria toxin-mediated ablation and chemogenetic activation of Mrgprb4-lineage neurons. 8-14 week old cage mate females (from different litters but shared cage for at least three weeks prior to testing) were acclimated to the behavioral room and brief experimenter handling for 1 hour the day prior to testing. Immediately prior to testing, the mice were separated in clean cages for 30 minutes to promote social interaction upon reunion. For chemogenetic experiments, the mice were injected with either saline or 0.5 mg/kg CNO in saline immediately prior to the 30 minute separation so that behaviors would be recorded 40-60 minutes after injection, when CNO has peak effects. The two females were reunited in the home cage for 30 minutes, during which video was recorded. The first 10 minutes served as habituation, during which time the mice predominantly explored the cage, and the following 20 minutes were scored for behaviors. Experimenter, blinded to genotype/treatment group, scored videos for duration allogrooming, duration selfgrooming, duration nestmaking, duration sleeping, number of conspecific crawl behaviors, and number of contacts received to the back. Behavioral observation research interactive software (BORIS) was used for behavioral scoring.

### Viral injections and optic fiber implantation

7-10 week old females were pretreated with 5mg/kg oral or intraperitoneal meloxicam before surgery. They were anesthetized in a chamber with 3% isoflurane before being placed in a stereotaxic frame and kept anesthetized with 1.5-2% isoflurane. Skull was exposed and leveled before drilling at the appropriate coordinates. 200nL pAAV-hSyn-GRAB_DA1h (Addgene) was injected unilaterally into NAc (AP: +1.0 mm relative to Bregma, ML: ±1.2 mm relative to Bregma, DV: 4.6 mm from the brain surface) using a backfilled glass needle and syringe pump (PHD Ultra, Harvard Apparatus). Skin was either sutured for one week recovery before implanting optic fiber (Doric, MFC_200/230-0.57_6mm_MF1.25_FLT) 200µm above injection site, or optic fiber was inserted immediately following injection. Three skull screws were inserted in the skull to stabilize the implant. A small amount of Metabond cement (Parkell) was used to bond the fiber, skull, and skull screws. Dental cement was applied on top of dried Metabond to create a stable implant structure. Mice were given 1 week to recover from implant surgery and 2 weeks to recover from injection and implant combined surgeries before behavioral testing. Meloxicam was administered 24 hours after surgery during post-operative monitoring.

### Fiber Photometry

Zirconia sleeves (Doric) were used to connect the optic fiber implant to the patch cord. Signals were recorded using a real-time processor (RZ10X, TDT) and extracted in real time using Synapse software (TDT). A 465nm LED was used to excite the GRAB_DA1h_ while a 405nm LED was used to measure changes in fluorescence due to photobleaching and movement artifacts.

For fiber photometry during transdermal optogenetic activation experiments, mice were placed on the mesh platform in plastic chambers with a 7mm slot cut into the ceiling to allow the cord to connect from the head to the computer. After 30 second baseline recording, 10Hz pulsed sin wave at 35mW blue light was shined through the mesh platform to the vaginal area at a 2-3cm distance for 30 seconds.

For fiber photometry during sexual behavior, females were placed in male home cage 1 hour into the dark cycle for 10 minutes or 10 mounts, whichever occurred first. If the male did not mount in the first 5 minutes, the female was placed in another male cage, for up to three total males.

TDT folders were imported directly into Fiber photometry Modular Analysis Tool (pMAT) software for analysis(Bruno et al., 2021). DeltaF/F was plotted as a peri-event time histogram from −10s to 20s, where 0s is the start of optogenetic stimulation to the anogenital region. deltaF/F calculated by normalizing to the median of 5s baseline sampling window (−10 --5s), bin constant: 150s.

### Electrophysiology

Recordings were made from *Mrgprb4*^*Cre*^*;Rosa*^*ChR2/+*^ or *Mrgprb4*^*Cre*^*;Rosa*^*ChR2/ChR2*^ mice (female; age 3.9 ± 0.6 wks). Mice were anaesthetized with ketamine (100 mg/kg i.p), decapitated, and spinal cord (T10-L2) rapidly removed in ice-cold sucrose substituted artificial cerebrospinal fluid (sACSF) containing (in mM): 250 sucrose, 25 NaHCO_3_, 10 glucose, 2.5 KCl, 1 NaH_2_PO_4_, 6 MgCl_2_, and 1 CaCl_2_. Sagittal slices (200µm thick) were prepared using a vibrating microtome (Leica VT1200S). Slices were incubated for at least 1hr at 22-24°C in an interface chamber holding oxygenated ACSF containing (in mM): 118 NaCl, 25 NaHCO_3_, 10 glucose, 2.5 KCl, 1 NaH_2_PO_4_, 1 MgCl_2_, and 2.5 CaCl_2_.

Following incubation, slices were transferred to a recording chamber and continually superfused with ACSF bubbled with Carbogen (95% O_2_ and 5% CO_2_) to achieve a pH of 7.3-7.4. All recordings were made at room temperature (22-24°C) and neurons visualized using a Zeiss Axiocam 506 color camera. Recordings were acquired in voltage-clamp (holding potential −70mV) or current-clamp (−60mV). Patch pipettes (3-7 MΩ) were filled with a potassium gluconate-based internal solution containing (in mM): 135 C_6_H_11_KO_7_, 8 NaCl, 10 HEPES, 2 Mg_2_ATP, 0.3 Na_3_GTP, and 0.1 EGTA, pH 7.3 (with KOH). No liquid junction potential correction was made, although this value was calculated at 14.7 mV (22 °C). All data were amplified using a MultiClamp 700B amplifier, digitized online (sampled at 20 kHz, filtered at 5 kHz) using an Axon Digidata 1550B, and acquired using Clampex software. After obtaining the whole-cell recording configuration, series resistance, input resistance, and membrane capacitance were calculated (averaged response to −5mV step, 20 trials, holding potential −70mV). Photostimulation intensity was suprathreshold (24 mW), duration 1 ms (controlled by transistor-transistor logic pulses).

## Acknowledgments

We thank members of the Abdus-Saboor and Abraira labs for helpful discussion and comments on this manuscript. We thank Oliver Hobert, Richard Axel, and Charles Zuker for helpful comments on this work and manuscript. We thank Nitsan Goldstein and members of the Betley lab for assistance with fiber photometry set up. We thank David Barker for suggestions with pMAT software. We thank members of LM’s thesis committee, including Lori Flannagan-Cato and Gregory Corder for helpful suggestions on experiments. We thank Qin Liu for sharing *Mrgprb4*^*Cre*^; *Rosa*^*ChR2/ChR2*^ mice with us. We thank Janet Sinn-Hanlon for illustrations. LM and MS are supported by NIH NRSA grants from NINDS and NCCIH. IAS and lab members acknowledge support from startup funds provided by the University of Pennsylvania and Columbia University, National Institute of Health grant NIH/NIDCR R00-DE026807, and fellowships from the Rita Allen Foundation and Alfred P. Sloan Foundation. VEA, MB, and MG acknowledge support from startup funds from Rutgers University, Pew Charitable Trust, NIH R01 and K01 grants from NINDS, and Whitehall Foundation.

## Notes

### Competing Interest Statement

The authors have declared no competing interest.

